# Reconstituting organotypic 2D microtissue co-cultures via sequential stenciling

**DOI:** 10.64898/2026.04.01.715780

**Authors:** Kai Hirzel, Jasmin Cic, Stella Asmanidou, Nadja Schmohl, Roland E Kontermann, Satoshi Toda, Monilola A Olayioye, Andrew G Clark, Michael Heymann

**Affiliations:** Institute of Biomaterials and Biomolecular Systems, University of Stuttgart, 70569 Stuttgart, Germany; Institute of Cell Biology and Immunology, University of Stuttgart, 70569 Stuttgart, Germany; Stuttgart Research Center Systems Biology, University of Stuttgart, 70569 Stuttgart, Germany; University of Tübingen, Center for Personalized Medicine, 72076 Tübingen, Germany; Institute for Protein Research, The University of Osaka, Osaka 565-0871, Japan

**Keywords:** micropatterning, co-culture, microtissue, tumor microenvironment, synthetic morphogenesis, intestinal homeostasis

## Abstract

In mammalian organisms, native tissue function depends on precise spatial organization down to the cellular level. Reconstituting tissue architectures in 2D *in vitro* platforms can provide a means to study direct and indirect cell-cell interactions in a variety of tissue contexts while remaining compatible with high-throughput assays and high-resolution live imaging. We combine cost-effective stereolithography leveraging 3D printing with replica molding to stencil spatially defined, multicellular culture systems with sub-millimeter resolution onto planar substrates. The system is designed for ease of use, requires no complex fabrication setups and scales readily to 96-well plates. Sequential stencil application and removal under a biosafety cabinet enables controlled positioning of multiple cell types and supported the maturation of tissue assemblies. We demonstrate the utility of this stencil-based patterning strategy in three applications. First, we employ a combination of two circular stencils to recreate a structural feature characteristic for the tumor microenvironment of solid tumors: the encapsulation of colorectal cancer cells by cancer-associated fibroblasts. Resulting cell patternings reproduce native tissue dynamics of the densely packed tumor tissues, in which cancer-associated fibroblast cells actively compress the cancer cells and confer targeted therapy resistance. Second, we probe the synthetic, diffusible morphogen system synNotch in patterned cell patches, where GFP-releasing cells generate a ligand-dependent gradient. Third, we recapitulate the characteristic crypt-villus architecture of the mammalian intestine by patterning intestinal organoids within a stencil-restricted crypt region and allowing differentiating cells to collectively migrate along a designed villus axis. The presented strategy allows for rebuilding multicellular tissue architectures *in vitro* with biologically relevant spatial precision for high-throughput drug screenings and dissection of tissue-specific cellular interactions.

## 1 Introduction

Mammalian tissues and organs are shaped by highly coordinated assembly processes during organismal development. Cells self-organize during early embryogenesis into three-dimensional, hierarchical structures upon mechanical and biochemical constraints, as well as dynamic, reciprocal interactions with their environment [1]. A series of spatiotemporally timed events during development culminate in individual cells assembling into functional, multicellular tissues [2]. Engineering relevant cell culture models with the accurate spatial *in vivo* tissue organization and function remains a field of active research [3, 4]. Traditional *in vitro* monolayer cultures in two dimensions (2D) were widely adopted in biomedical research for their ease of use, cost-effectiveness and compatibility with high-throughput screening assays decades ago [5]. However, any flat cell sheet will ultimately fail to recapitulate the complete microphysiology of a native tissue [6, 7]. Consistently, drug candidates with promising preclinical results obtained from cultured 2D cell layers underperform too often in subsequent clinical patient trials [8]. An ideal *in vitro* model has a high predictive value to translate seamlessly into the clinic. For this, three-dimensional (3D) *in vitro* systems including spheroids, organoids [9, 10], organ-on-chip systems [11] and bioprinted tissues [12] have been actively developed, with each technology opting for different tradeoffs between desired utility against preparation complexity and hence cost and availability.

Multicellular arrangements in 2D can be readily implemented with only modest technical effort in routine cell biology laboratories and represent an attractive middle ground between reductionist 2D monolayer and complex 3D culture systems [13]. Multiple cell types may be seeded randomly [14, 15, 16] or by controlling their adhesion on the culture substrate to defined regions [17, 18]. Commonly used techniques to pattern living cells onto planar culture substrates include biochemical conjugation methods like click-chemistry or immunocapture [22, 23], selective adhesion [24, 25] and physical techniques such as mechanical constraints [26] or stamping [27, 28]. Elastomeric stamps or stencils fabricated via soft lithography are now routinely used for protein- and cell-deposition on 2D substrates for a variety of applications. For example, microcontact printing has been used to pattern human induced pluripotent stem cells for differentiation studies [29]. Further, microfluidic stamps of thin polydimethylsiloxane (PDMS) films have been applied to pattern multiple cell types in defined regions aligned with microfluidic channels [30].

During development, spatial tissue organization is guided by morphogen gradients [31]. Morphogens are diffusible molecules that allow cells to interpret their relative location within a tissue and provide cues that guide cell differentiation and cell fate decisions [32, 33]. A central aim in synthetic biology is to reconstitute gradient-driven developmental processes *in vitro*. The synthetic Notch (synNotch) system exploits chimeric forms of the Notch transmembrane receptor, in which both the extracellular binding domain and the intracellular signal-transmitting domain are replaced with custom designed, heterologous protein modules [34]. Upon ligand binding, the engineered receptors undergo a conformational change, releasing the intracellular domain into the nucleus to activate target gene transcription. In this way, orthogonal signaling circuits can be constructed that act independent of endogenous pathways. SynNotch signaling cascades have been applied to multi-cellular patterning and self-organization into multidomain structures of mammalian cells [35, 36]. Tethering a diffusive molecule such as green-fluorescent protein (GFP) ligand to an anti-GFP synNotch receptor was used to generate positive and negative feedback loop mediated signaling patterns.

A central example of how spatial organization of multiple cell types is critical for specialized tissue function is the mammalian intestine. The small intestine is lined by a single-cell sheet of polarized epithelial cells that are compartmentalized into crypt and villus domains. Crypts are inward-folding structures that protect stem cells from the luminal environment and protruding structures, called villi, that project into the intestinal lumen [38]. Intestinal stem cells in the crypts undergo constant self renewal and give rise to rapidly dividing transit amplifying cells, which further differentiate into absorptive enterocytes, as well as various secretory cell types. Differentiated cells migrate to the tip of the villi, where they perform nutrient absorption and secretion before being extruded from the monolayer [39]. This zonal organization protects stem cells while maximizing surface area, enhancing absorptive capacity to absorb digested nutrients [40]. This architecture has been partially recapitulated *in vitro* using organoid cultures derived from intestinal stem cells, which self-organize into 3D structures containing crypt-like and villus-like domains and can maintain proliferation and differentiation programs reminiscent of the native tissue [41, 42]. However, intestinal organoids suffer from limited experimental reproducibility and poor control over spatial gradients, and their enclosed 3D geometry makes cell-scale imaging challenging [43, 48]. Several gut-on-chip model systems have succeeded in reproducing aspects of the 3D crypt-villus architecture with improved microenvironmental control, yet these platforms remain technically demanding and similarly pose difficulties for high-resolution imaging [49, 50, 51]. Alternatively, organoid-derived monolayers offer improved accessibility but lack defined spatial organization, where randomly distributed crypts lead to chaotic migration patterns of differentiated cells around the crypts.

Multicellular organization can also be observed as a defining feature in diseased tissues, such as the cancerous tissue of solid tumors, for instance in colorectal cancer (CRC). Here, cancer cells coexist and associate with native healthy cells forming the heterogenous tumor microenvironment (TME) [52, 53]. Within the TME, cancer-associated fibroblast (CAF) cells have been recognized as essential to remodel the extracellular matrix within the tumor tissue. These CAF cells often align around cancer cells, compressing them by exerting mechanical forces [54, 55]. In colorectal cancer, CAF cells act as functional drivers of tumor progression by providing biochemical and mechanical cues that promote cancer-typical hallmarks such as angiogenesis, immunomodulation, and cancer cell invasion [56, 57]. In therapeutic treatment regimes, CAF cells can confer resistance to targeted anti-cancer drugs by secreting survival factors and by acting as a physical barrier that prevents soluble suppressive drugs from reaching the cancer cells [55, 58]. Although two-dimensional co-cultures of CRC and CAF cells have become a common alternative to CRC cell monocultures, spatial organization of both cell types is difficult to control when seeding randomly. Conventional cell monocultures often fail to reconstitute a functional tumor-stroma interface and cancer cell invasion dynamics [59]. Furthermore, cancer cells can exhibit distinct behaviors within the TME core compared to the tumor periphery. This structural arrangement of tumors is absent in randomly seeded co-cultures [60].

Despite the importance of multicellular organization for tissue function, broadly accessible *in vitro* model systems that are capable of reflecting how spatial arrangements shape cellular behaviour remain scarce in life science laboratories. To address this, we patterned 2D multicellular micro-tissues using custom replica-molded PDMS stencils (Figure 1). Stencils were fabricated by soft lithography using 3D-printed master molds that enabled multiple rounds of stencil production without the need for complex microfabrication infrastructure or cleanroom facilities. We optimized the fabrication process to achieve cell patterning at physiologically relevant scales of below 1 mm. Next, we tuned the PDMS curing time for the stencils to adhere reversibly to standard cell culture substrates. This allowed us to restrict initial cell attachment as well as long-term cell and organoid cultivation within defined zones. Using this approach, we were able to successfully design and implement (1) a 2D reconstruction of the tumor microenvironment using spatially organized colorectal cancer cell and fibroblast co-cultures, which reproduced native CAF-driven compression and drug resistance, (2) linear and radial morphogen diffusion gradients using the synNotch system and (3) a recapitulaition of the intestinal crypt–villus architecture, enabling directed cell migration along the villus region. Sequential insertion and/or removal of multiple stencils enabled the patterning multiple cell types and organoid assemblies in a controlled geometric arrangement to mimic naturally architectures of multiple cell types.

**Figure 1:**
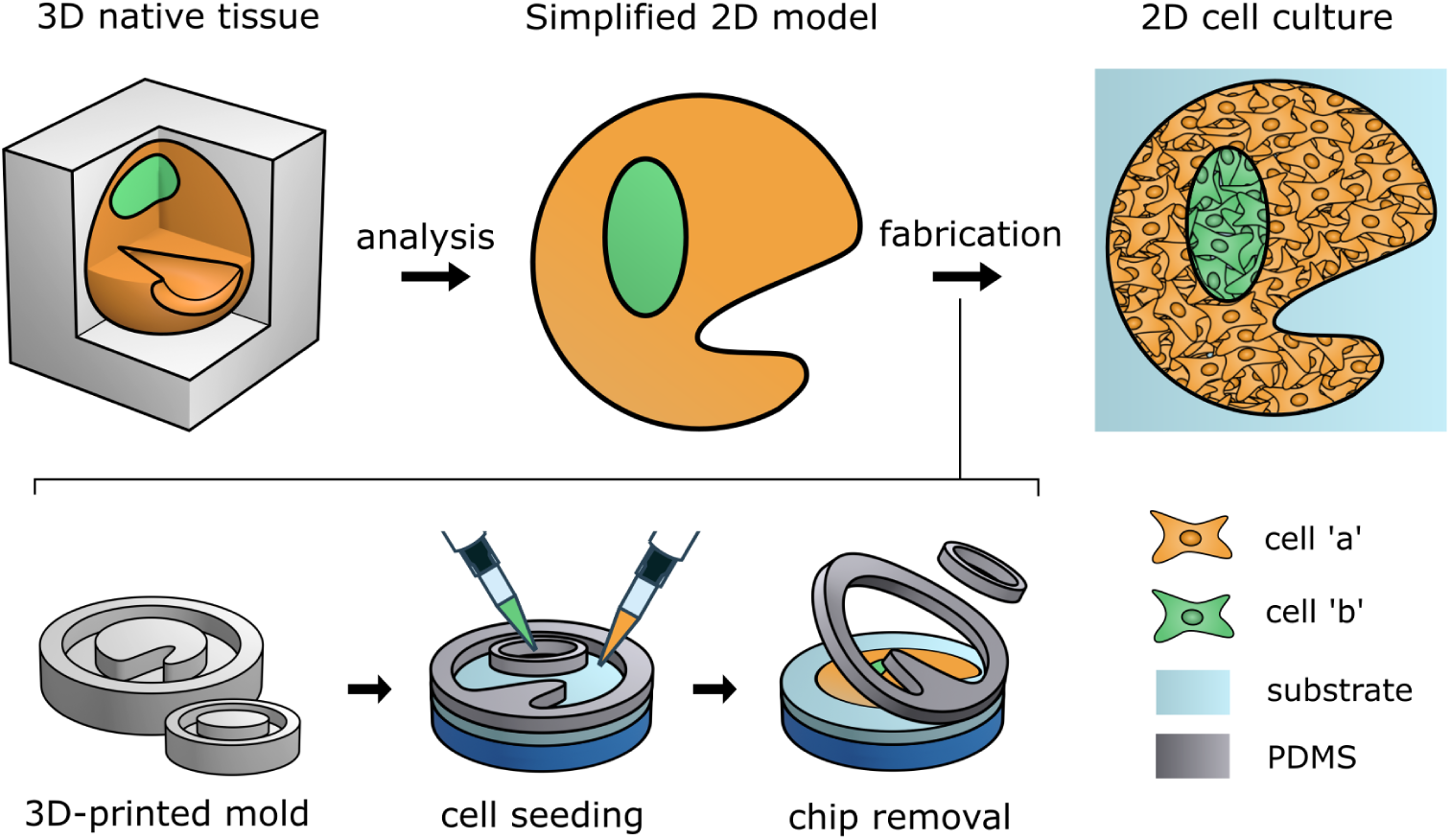
Projecting physiological 3D tissue architecture into a 2D multicellular pattern model. Representative *in vivo* morphologies and cellular architectures of 3D native tissue inform a simplified 2D model for multicell patterning. While not replicating all details, the 2D projection seeks to preserve essential features in an experimentally tractable *in vitro* format. Here, two distinct cell types ‘a’ and ‘b’ are patterned. 3D-printed molds of the desired geometry are replica molded into PDMS stencils. The stencils can transiently attach to standard cell culture substrates to restrict the local cell adhesion area, generating confluent multicellular patches of the target geometry.

## 2 Results

### 2.1 3D print resolution for optimal replica molding

Soft-lithography of microstencils typically involves high-resolution master molds produced with specialized and complex microfabrication strategies such as photomask lithography or two-photon 3D printing [61, 62]. In contrast, one-photon stereolithography (SLA) 3D printing offers a low-cost, rapid and infrastructure-free alternative for master mold fabrication at the expense of the minimally achievable feature resolution [63]. To fabricate master molds for soft lithography, we first evaluated the print accuracy of the used SLA 3D printer. Print resolution is defined by the minimal volume element (voxel) that can be achieved. Illuminated voxel dimensions are defined by the lateral focal spot size specified for the instrument and the axial penetration depth into the print resin [64]. Resulting written voxel dimensions can differ substantially however, as print part orientation (relative writing direction), photochemical resin properties or exposure time can each affect voxel polymerization, and hence overall dimensions [65, 66]. A gold-standard to access lateral resolution is described by the Rayleigh two-point criterion [67], where the midpoint intensity between two identical light points is defined as:

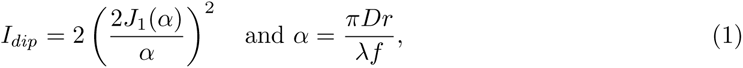

when two diffraction-limited points are just resolvable, as the peak of the first diffraction pattern (or Airy disk) aligns with the first minimum of the second [68]. In optical systems limited by diffraction, the point spread function of a circular aperture is described by the Airy pattern with *J*_1_(*α*) as the first-order Bessel function, *α* as the scaled radial coordinate, *D* as the aperture diameter, *λ* as the wavelength of light, *f* as the focal length of the imaging system, and *r* as the radial distance from the center of the Airy disk [69]. For this, *I_dip_* is quantified as *I_max_/I_min_* from a line profile view in lateral direction. At the Rayleigh limit, the combined intensity at the midpoint (dip) halfway between both points results in: *I_dip_ ≈* 2 *·* (0.6076)^2^ = 0.7384. Hence, a dip to approximately 74% of the peak intensity is still accepted as resolvable.

While numerous target designs have been explored to assess optical resolution [70, 71], the 1951 US Air Force (USAF) target remains a widely adopted benchmark to define the Rayleigh criterion [72]. To assess the smallest resolvable feature for replica molding, the USAF target was extruded to a height of 200 µm and 3D-printed (Figure 2A, design 1). The largest line pair (group -2, element 1) had a line width (*lw*) = line distance (*ld*) = 2 mm and a line length (*ll*) = 10 mm. 3D-printed USAF targets were imaged by brightfield microscopy, fluorescently stained, and subsequently analyzed in 3D from z-stack images obtained by multiphoton microscopy. Print lines deviated consistently by about +100 µm in both lateral direction, corresponding to an overcuring of each edge by about 50 µm (Figure 2A). With an *I*_dip_ = 0.75 lines down to group 1, element 2 still satisfy the Rayleigh criterion, but are ill suited for replica molding as lines pairs overlap too strongly (Figure 2B). We thus define a 50% line-pair flatness (lpf) as our resolution criterion for replica molding, as the line pair that still maintained flat for at least half the nominal line width (Figure 2B). This 50% in-line flatness criterion was met down to group 0, element 3, or a nominal lines spacing of 400 µm.

**Figure 2:**
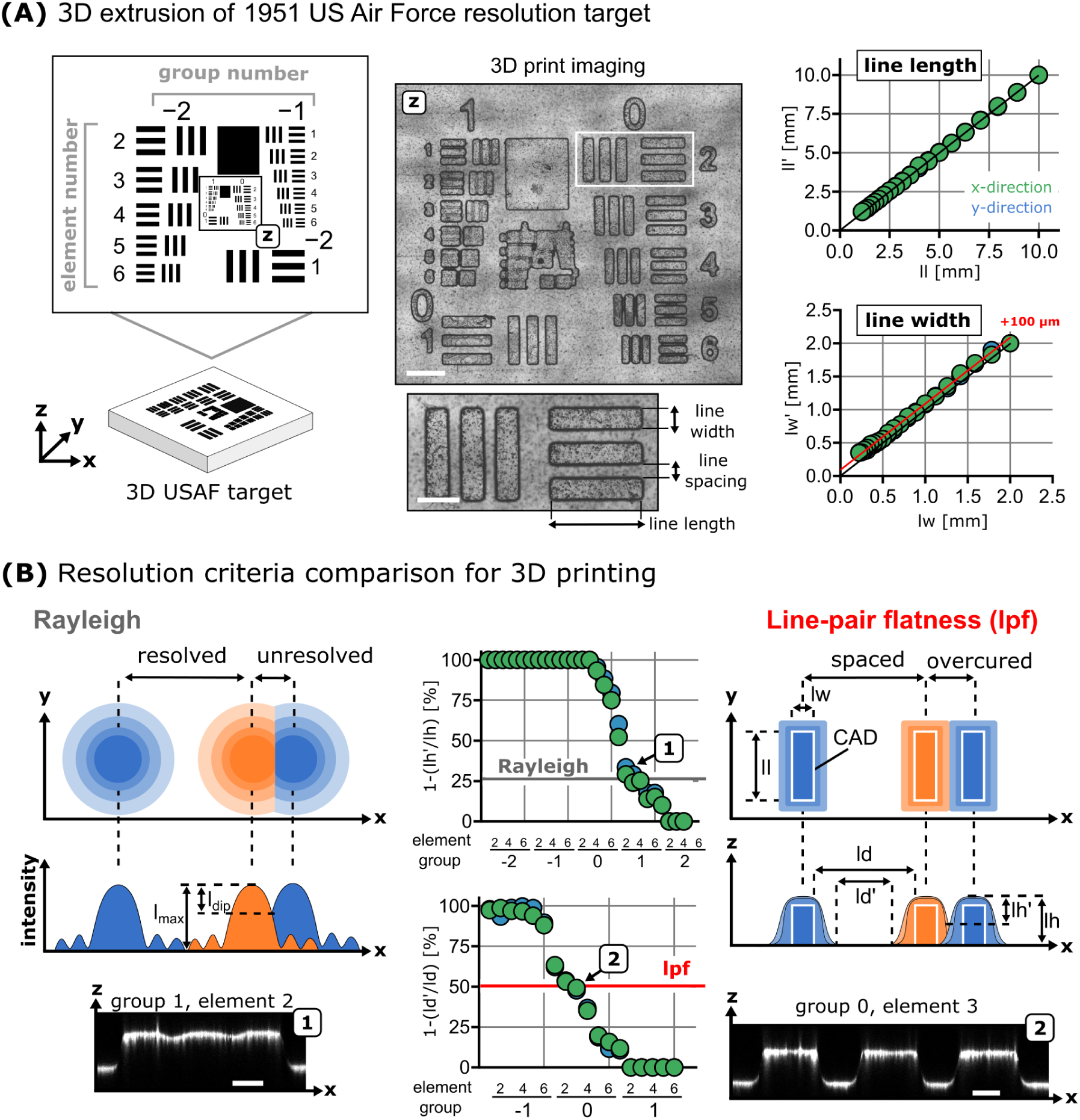
Metrology of the 3D printing process to identify optimal feature sizes. **(A)** Extruded 3D USAF target design to assess the smallest resolvable feature for replica molding. Brightfield imaging reveals merged line pairs for higher group elements (scalebar 2 mm, zoom-in 1 mm). Printed line width and line length exceed the nominal CAD dimensions by about 100 *µ*m. **(B)** The Rayleigh resolution limit defines two points as yet resolved, if their overlap zone does not exceed about 74% of the peak intensity. For the printed lines pairs this condition is still satisfied down to group 1, element 2, corresponding to a line width of 200 *µ*m. The more stringent line-pair flatness criterion requires a flat surface for at least half the space between two neighboring lines, which was satisfied down to group 0, element 3, or about 400 µm gap between both lines (scale bars 200 *µ*m).

### 2.2 PDMS stencil fabrication via soft lithography and cell patterning

To achieve leakage-free cell patterning, stencils must adhere firmly on a planar substrate yet be able to be removed cleanly from it. For this, we sought to identify curing conditions that yield PDMS with the appropriate balance of elasticity and tackiness (Figure 3A). Wear resistance, elastic modulus, and yield stress of PDMS elastomers result from the balance between cross-linkable monomers and reactive linker and hence overall cross-linking density, the degree of polymer crystallization, and the curing conditions, including temperature and curing time [73]. In a first step, curing conditions were optimized to yield flexible and tacky PDMS stencils (Figure 3B). A low curing agent to monomor ratio of 1:30 failed to cure fully, and uncured PDMS debris remained on the glass substrate during attach-detach cycles. More commonly used ratios between 1:10 and 1:15 showed weak surface adhesiveness and were hence deemed overcured. Finally, 45 minute curing of a 1:20 ratio at 80 °C achieved leakage-free incubation up to 7 days and could also be removed without leaving residual staining on the substrate (Figure 3B).

**Figure 3:**
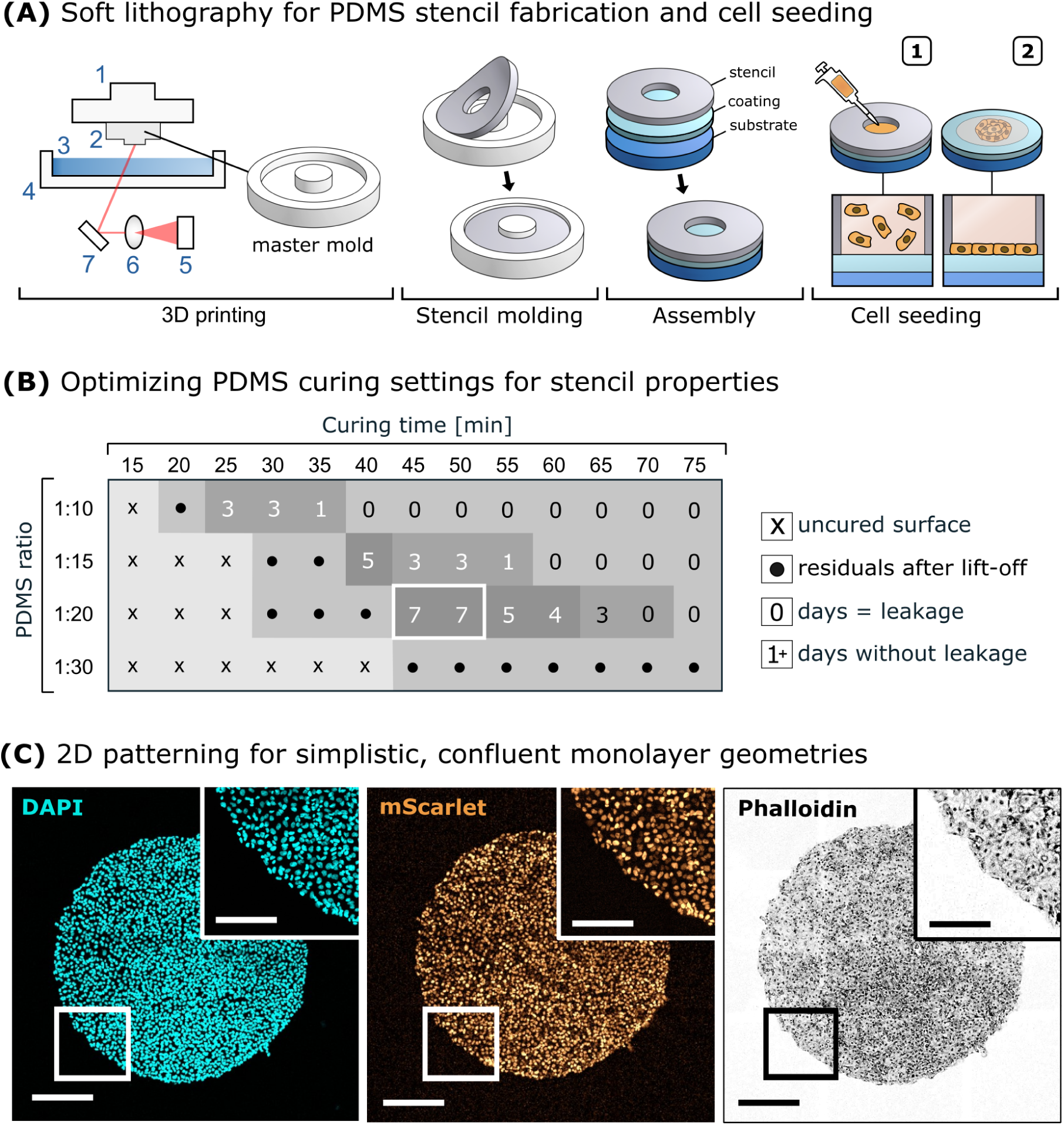
Micromold fabrication for cell patterning. **(A)** Schematic workflow for stencil fabrication and cell loading. Master molds were 3D-printed using 1PP SLA to stencil PDMS (1 = print head, 2 = print structure, 3 = print resin, 4 = resin tank, 5 = laser source, 6 = optical lens, 7 = mirror device). Attached stencils were loaded with the corresponding cell solution (1), cells patterned and cultivated within the restricted adhesion area inside the stencil channels (2). **(B)** Stencils were evaluated for shortened curing times and PDMS to curing agent ratios for optimal surface ‘tackiness.’ A 1:20 ratio with curing for 45 minutes cured at 80 °C provided optimal substrate attachment without leakage for 7 days. **(C)** Confocal imaging confirmed sharp circular cell patches with intact confluency and precise contour replication after stencil removal (scale bar 250 *µ*m, zoom-in scale bar 100 *µ*m).

We next sought to optimize the stencil patterning process for cell deposition. For ease of use under the cell culture bench, a 1 mm diameter, circular cell monolayer, referred to as a cell patch, was chosen, as this could easily resolved by eye without additional magnification while also still being near physiologically relevant dimensions (design 2). Stencils were placed with tweezers onto a round collagen-coated cover slide. Then, a 2 µl cell solution containing 1,000 colorectal LIM1215-mScarlet cancer cells was loaded into the stencil opening with a 10 µl pipette. Within 24 hours, cells adhered and formed a confluent monolayer, at which time stencils were removed (Figure 3C). Fluorescent phalloidin staining revealed that actin filaments in resulting cell patches followed a sharp circular contour matching the stencil rim. Also, stencil removal did not disrupt the established pattern, suggesting that peripheral cells did not adhere to the stencil during cultivation.

Finally, PDMS stenciling should not adversely affect cultured cells. While generally considered biocompatible, small molecules can diffuse into and out of the PDMS polymer matrix [75]. For example, released PDMS mono- and oligomers have been shown to absorb into cells and to interfere with their metabolism [76]. We thus assessed the effect of stencils on colorectal LIM1215-mScarlet cancer cells and on primary, immortalized cancer-associated IKP-CAF-001-GFP fibroblast cells (SI figure 1). Cells were seeded on a collagen-coated cover glass slides (1) with no further modification as a control, or (2) confined into a 1 mm stencil. Alternatively, a closed PDMS stencil was pre-incubated for 24 hours on the collagen-coated cover glass surface before seeding cells (3) without further confinement, or (4) confined into a 1 mm stencil. Cellular substrate attachment and cell spreading was unaffected for all four conditions after 24 hours of cultivation (SI figure 1). Also, cellular proliferation remained similar for at least 72 hours of cultivation. Cellular ATP levels confirmed consistent metabolic activity 24 hours after seeding/-patterning across all tested conditions, with all relative ATP level falling within *>*97% of the normalized control.

### 2.3 Patterning cancer-associated fibroblasts as a tumor microenvironment model for drug screening

CAFs are associated with therapeutic resistance and tumor progression [77]. Developing targeted tumor treatments would benefit from suitable experimental models of the CRC-CAF microenvironment. To recapitulate the radial CRC enclosure by CAF cells, we combined an inner (S_in_, design 3) and outer stencil (S_out_, design 4) (Figure 4A). As model cells, we used the CRC cell line LIM1215 stably expressing the fluorescent protein mScarlet and an immortalized CAF cell line with stable GFP expression. Stencils were removed 24 hours after seeding, leaving an approximately 1 mm gap between the 1 mm diameter central CRC and surrounding CAF patch. Both cell monolayers started to proliferate towards each other, whereby they continuously covered the previously restricted zone. Within 24 hours, both patches fused to a closed cell layer with a well defined CRC-CAF interface (Figure 4B). This interface showed increased cell densities when compared to the initial stenciled contours. Also, individual LIM1215-mScarlet cells began migrating into the CAF area after reaching full confluency, indicating that CRC cell migration was not inhibited by the presence of the CAF area. During subsequent incubation the central LIM1215-mScarlet patch decreased by 20% over 24 hours, consistent with a fibroblast mediated compression force against enclosed CRC cells [55] (Figure 4C). This was not the case in a control experiment, where mScarlet expressing LIM1215 cells readily intermixed with their surrounding non-fluorescing LIM1215 counterparts, resulting in an increase of the central patch by 30% and a rather unlabeled contact line between both zones 24 hours after reaching confluency.

**Figure 4:**
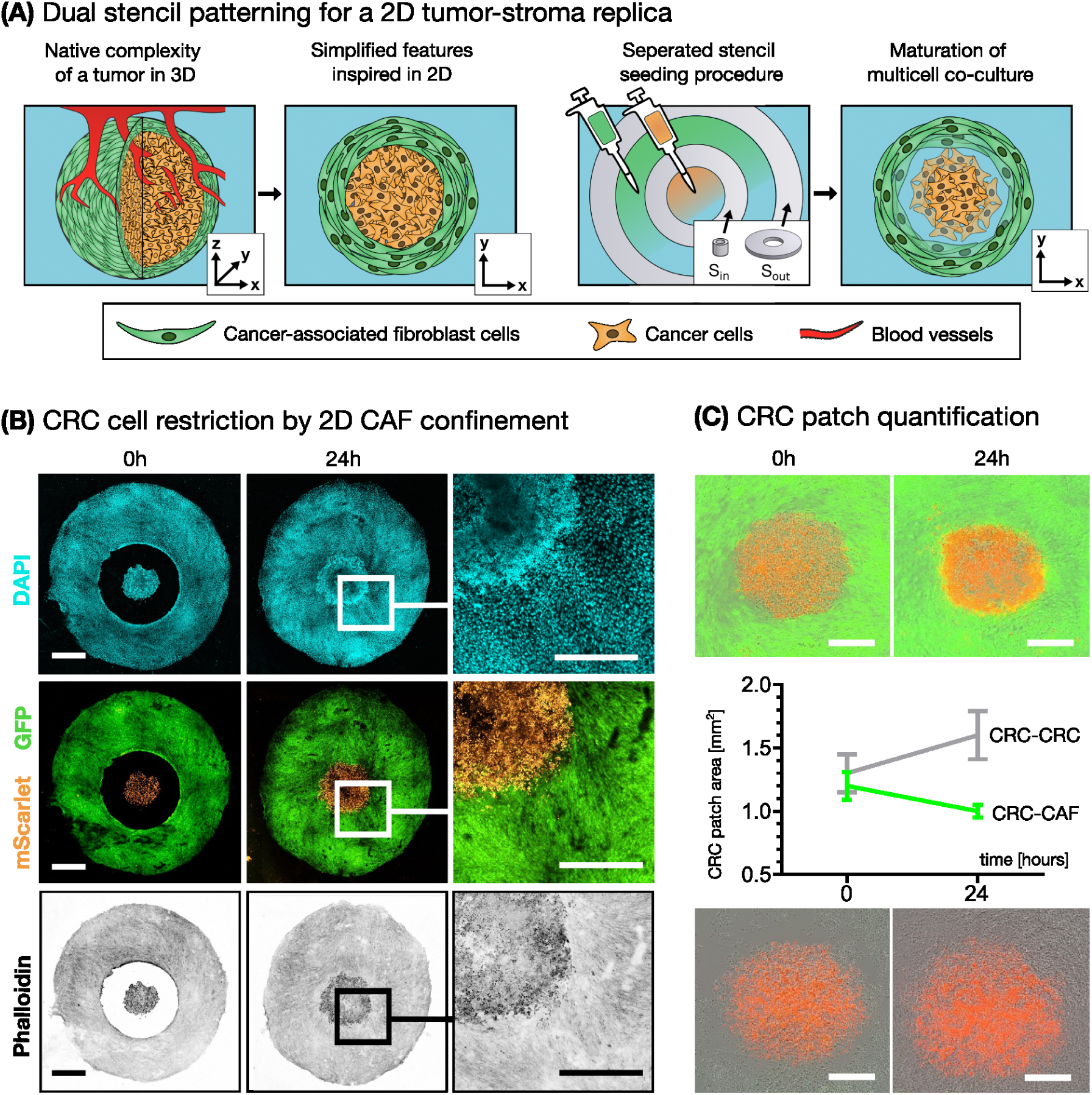
Dual stencil approach to pattern cancer and cancer-associated fibroblast co-cultures. **(A)** Process schematic for the concentric enclosure of CRC cells by CAFs. Two circular stencils were placed into each other with S_in_ in the center of the stencil channel of S_out_. Loading each stencil separately yields two monolayer patches that grow together after removal of S_in_. **(B)** Confocal imaging showed stencil gap closure by LIM1215-mScarlet and fibroblast cells (IKP-CAF-001-GFP) within 24 hours after removal of S_in_ (scale bar 1 mm, zoom-in scale bar 500 *µ*m). **(C)** Live-cell imaging revealed a CAF-induced compression effect (scale bar 500 *µ*m). CRC patch area strongly declined in CRC-CAF, but increased in CRC-CRC co-cultures 24 hours after stencil gap closure. CRC patch area was quantified by ROI image segmentation based on the mScarlet fluorescence signal (n=3, N=15, error bars show the mean *±* SD).

To investigate whether drug response within the colorectal cancer cells is influenced by heterotpyic interactions between cancer cells and the CAF cells, we next applied cetuximab and afatinib to CRC-CAF and to CRC-CRC control co-cultures. The monoclonal antibody cetuximab is commonly administered against metastatic CRC. Cetuximab blocks EGFR activation to suppress a HER family receptor tyrosine kinase that is overexpressed in over 60% of colorectal tumors [78]. However, a high abundance of CAF cells in a tumor dilutes EGFR suppression by cetuximab, as the antibody can be sequestered by the fibroblasts. Furthermore, cetuximab stimulates HER-ligand secretion including EGF from CAF cells, which confers resistance to Cetuximab by activating EGFR signaling [58]. The small molecule tyrosine kinase inhibitor afatinib can counteract this compensatory signaling mechanism by irreversibly blocking the kinase activity of EGFR, HER2, and HER4 [79].

In order to determine how the presence of CAFs influences therapeutic efficacy, we treated CRC-CAF and CRC-CRC control co-cultures with cetuximab and afatinib and observed by live-cell imaging over 72 hours. The LIM1215-mScarlet cancer cell patch area was quantified with ROI image segmentation based on the mCherry fluorescence signal (Figure 5). In PBS and DMSO controls, both co-cultures showed a continuous increase of the respective CRC patch sizes. Cetuximab treatment reduced CRC patch growth by about 50-60% in CRC-CRC co-cultures but had no significant effect on CRC-CAF co-cultures, which is consistent with a CAF mediated cetuximab resistance of CRC cells. In contrast, afatinib inhibited CRC patch growth with and without CAF co-culture by 50-70%. Thus, cetuximab inhibited CRC cancer cell patch growth only in the absence of CAFs, whereas afatinib suppressed CRC cancer cell patch expansion in both CRC-CAF and CRC-CRC co-cultures, consistent with afatinibs capability of a broader blockade of multiple HER family receptors. Taken together, the co-culture patterning approach could hence recapitulate drug treatment responses consistent with clinical observations and provides a promising alternative to conventional cancer cell monocultures for drug screenings.

**Figure 5:**
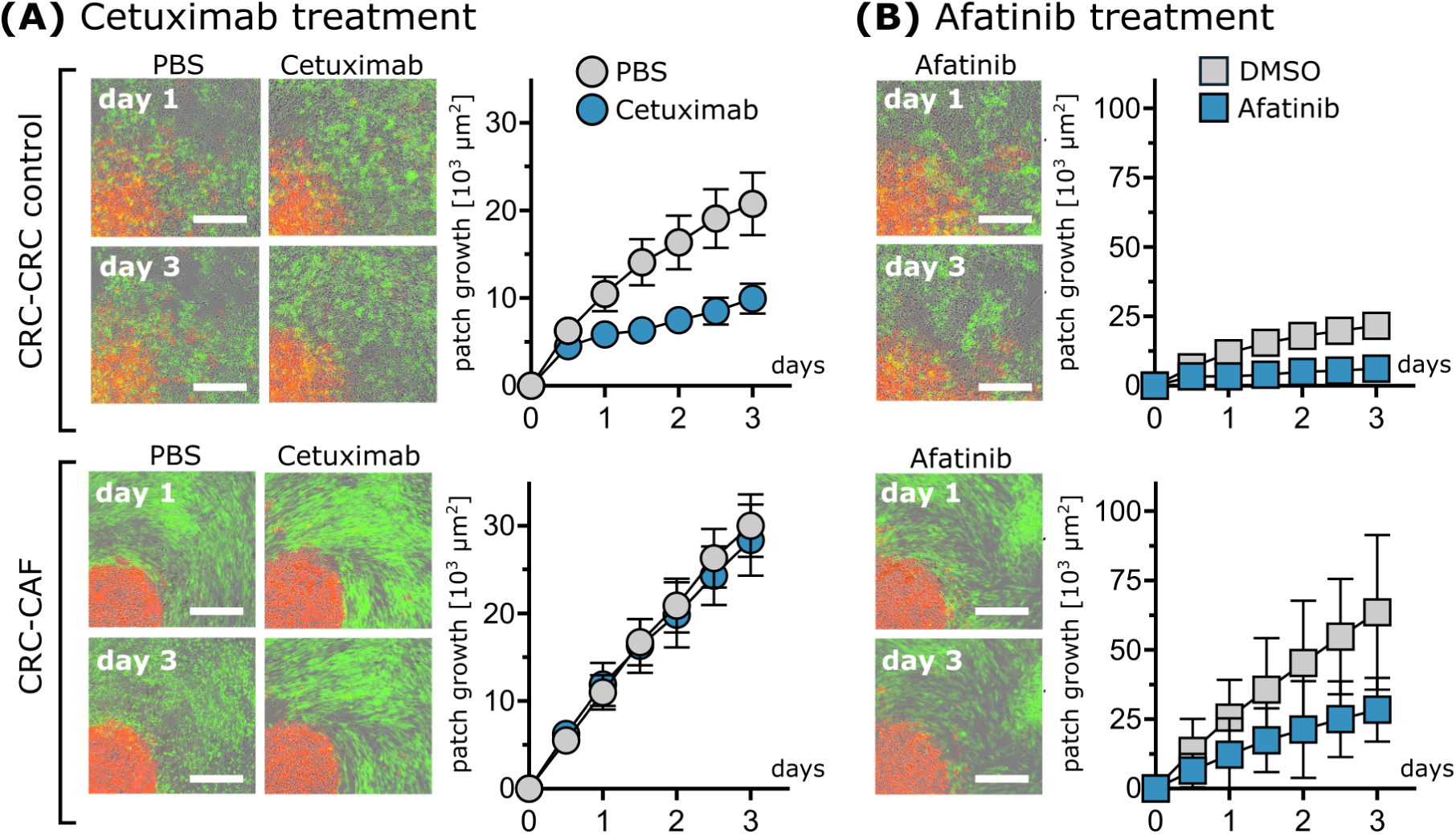
LIM1215-mScarlet drug response in CRC-CRC and CRC-CAF co-cultures over 72 hours of incubation. CRC patch area was quantified by ROI image segmentation based on the mScarlet fluorescence signal (n=3, N=30, error bars show the mean *±* SD). **(A)** Cetuximab treatment resulted in overall smaller CRC cell patches in CRC-CRC co-cultures, compared to a drug free control. In CRC-CAF co-cultures in turn showed no significant difference between treated and untreated control over three days, indicating a CAF-mediated protection. **(B)** CRC patch size in both the CRC and CAF background exhibit a smaller show reduced growth under afatinib treatment compared to untreated controls.

### 2.4 Patterning synthetic morphogenic signaling gradients

After reconstructing a native, pathology-relevant tissue interaction, we next asked whether the cell patterning strategy could also be applied to evoke signaling patterns within a synthetic system. Self-organizing cell patterns can be generated using a synNotch-based morphogen reaction diffusion system [36]. This system consists of sender cells that secrete GFP, receiver cells that express a synNotch receptor engineered to bind and be activated by GFP and an anchor protein that catches and presents GFP to the synNotch receptor (Figure 6A). GFP binding to receiver cells induces an intracellular cleavage of the synNotch receptor, such that its intracellular domain can enter the nucleus to activate mCherry expression. In this system, GFP acts as a morphogen that can activate nearby receiver cells, but becomes depleted at higher distances from a sender cell pole. Resulting patterns strongly depend on the specific details of the reaction circuit, but also on the specific geometry.

**Figure 6:**
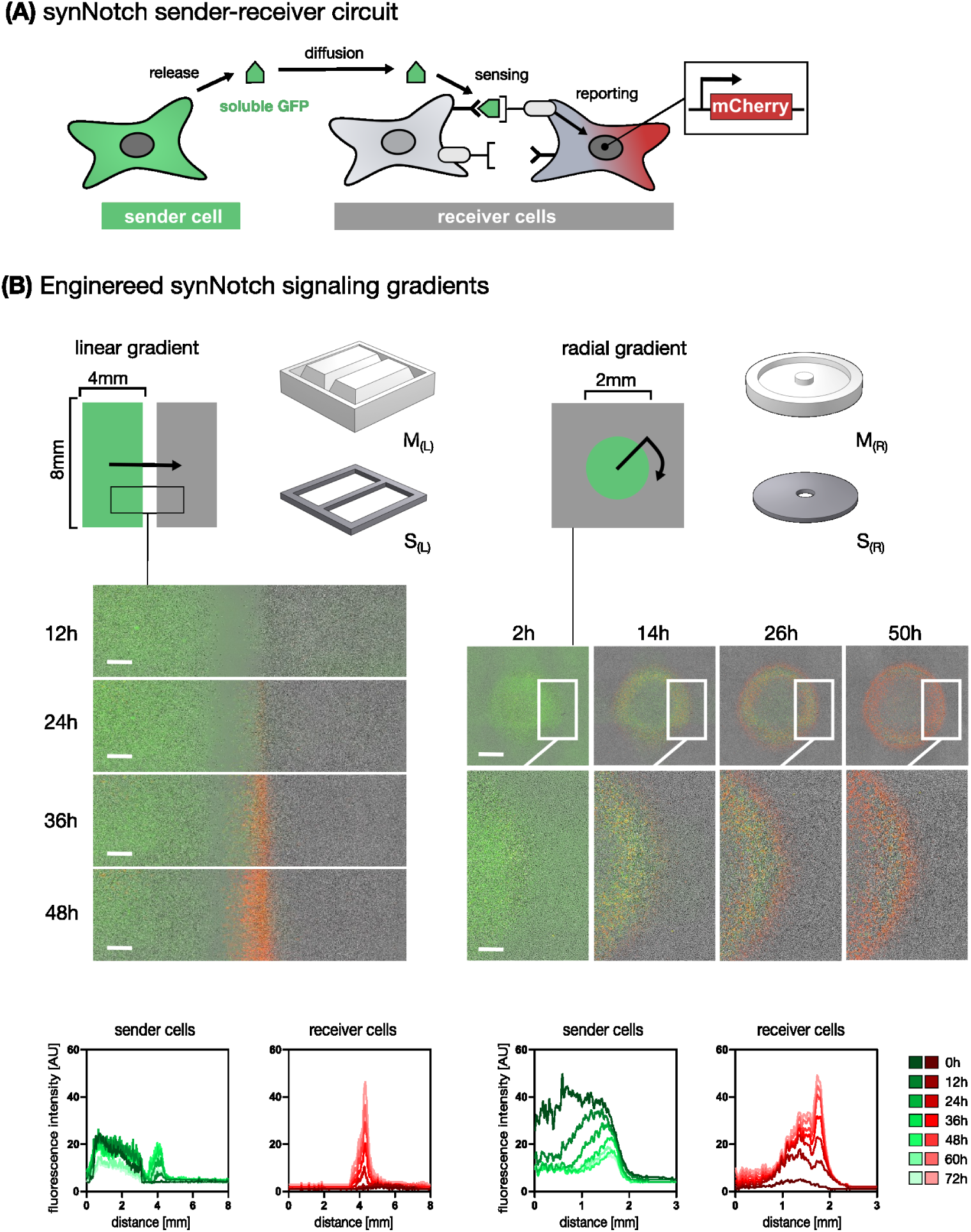
Patterned co-cultures to engineer synthetic signaling gradients. **(A)** Schematic of synNotch sender-receiver circuit. Sender cells secrete soluble GFP, which diffuses and is captured by anchor cells to induce synNotch mediated mCherry production in neighboring receiver cells. **(B)** Organizer centers were designed as rectangular or circular sender cell patches that couple into a confluent monolayer of receiver cells to yield linear or radial diffusion patterns respectively. Live cell imaging revealed the formation of an mCherry fluorescence gradient (time points after cell seeding, scale bar 250 *µ*m).

Sculpting organizer centers as sources or sinks for morphogen gradients, as well as their size and relative position can help to advance our understanding of the self-organization and pattern formation in biological systems. We hence stenciled GFP sender cells either into a 4 mm x 8 mm rectangle with a neighboring tissue domain of a monolayer of receiver cells (S_L_, design 5) or into a round 2 mm radius pole (S_R_, design 6) that was overlaid and surrounded by a monolayer of receiver cells (Figure 6B). GFP morphogen gradients stabilized within 48 hours, while a characteristic mCherry fluorescence reporting receiver cell activation emerged within 24 hours. mCherry fluorescence activation followed a clear gradient prescribed from the initially formed GFP morphogen gradients and generated linear or radial gradients depending on the geometry of the sender-receiver regions, confirming the utility of the experimental cell stencil protocol.

### 2.5 Intestinal organoid micro-tissue patterning of the crypt-villus axis

Conventional 2D cell and organoid monolayers typically display random cell alignment and hence do not sufficiently replicate the precise spatial organization characteristic of the mammalian intestine. In the intestine, stem and progenitor cells located in crypts divide and then differentiate while migrating toward the villus tip [80]. Intestinal organoids and micro-physiological systems show self-renewal and lineage differentiation *in vitro*, but are labor-intensive and scale poorly [81, 82, 42, 43]. By contrast, 2D organoid monolayers preserve key epithelial features as polarity and barrier function, while supporting high-resolution imaging [41, 83].

To construct a spatially functional mimic of the intestinal epithelium, we used a dual stencil configuration for patterning intestinal organoids in a stencil-restricted crypt zone in 2.5D. First, a crypt-villus stencil (S_cv_, design 7) was placed on an L-DOPA-coated polyacrylamide hydrogel. Second, a solution of collagen-I and laminin 1 was deposited into the stencil opening to funcitonalize the entire crypt-villus pattern. Next, a villus-blocking stencil (S_block_, design 8) was used to obstruct the villus section of the first stencil. Villi were sized to be 0.8 mm long to match a typical human intestinal villus height of about 0.4 - 1.0 mm [84]. A vernier scale was included with the stencil design to achieve 50 µm alignment precision when working without magnifying optics in the clean bench (Figure 7) [44]. Intestinal organoids were then seeded into the crypt zone to form an initial monolayer, from which differentiated cells started to migrate toward the villus within 24 hours after removing both stencils (S_block_) (Figure 8A). Following stencil removal, crypt-like regions of the organoids remained stationary in the crypt compartment, while differentiated cells migrated into the villus compartment, forming a confluent monolayer throughout the entire crypt-villus pattern. Particle image velocimetry (PIV) analysis revealed directed migration along the villus axis, suggesting that differentiated cells established collective migration streams even after full confluence had been reached (Figure 8B). Migration velocities along the crypt-villus axis peaked at 2 *µ*m/h and decreased towards the tip of the villus, similar to migration dynamics from *in vivo* studies and live imaging of intestinal explants [45]. Control samples without S_block_ displayed random migration patterns along crypt and villus. Taken together, the synthetic crypt-villus axis demonstrates how the sequential stenciling approach can be used to spatially pattern primary organoids to specific geometries and reproduce native tissue dynamics.

**Figure 7:**
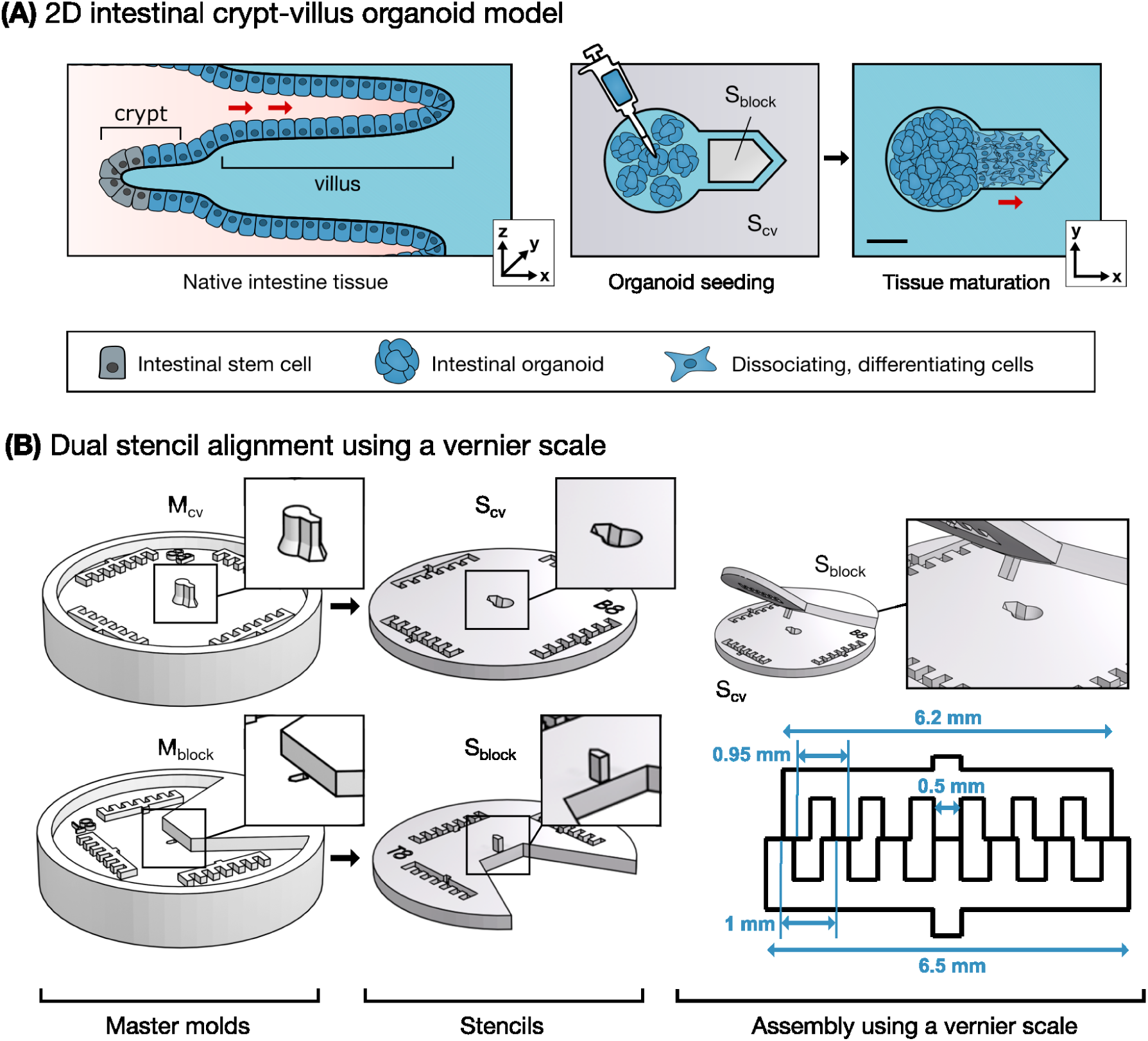
Patterning of a 2D intestinal crypt-villus axis. **(A)** The characteristic crypt-villus topography was projected into a circular crypt region with an extruding villus zone for migrating and differentiating cells. Intestinal organoids were seeded within the crypt of S_cv_ while S_block_ closed the crypt transiently. The red arrow indicates the path for collective migration of differentiating cells along the villus axis (scale bar 500 *µ*m). **(B)** S_cv_ and S_block_ included a vernier scale for precise stencil alignment by eye to help position S_block_ in the center of the villus to restrict organoid seeding in this zone.

**Figure 8:**
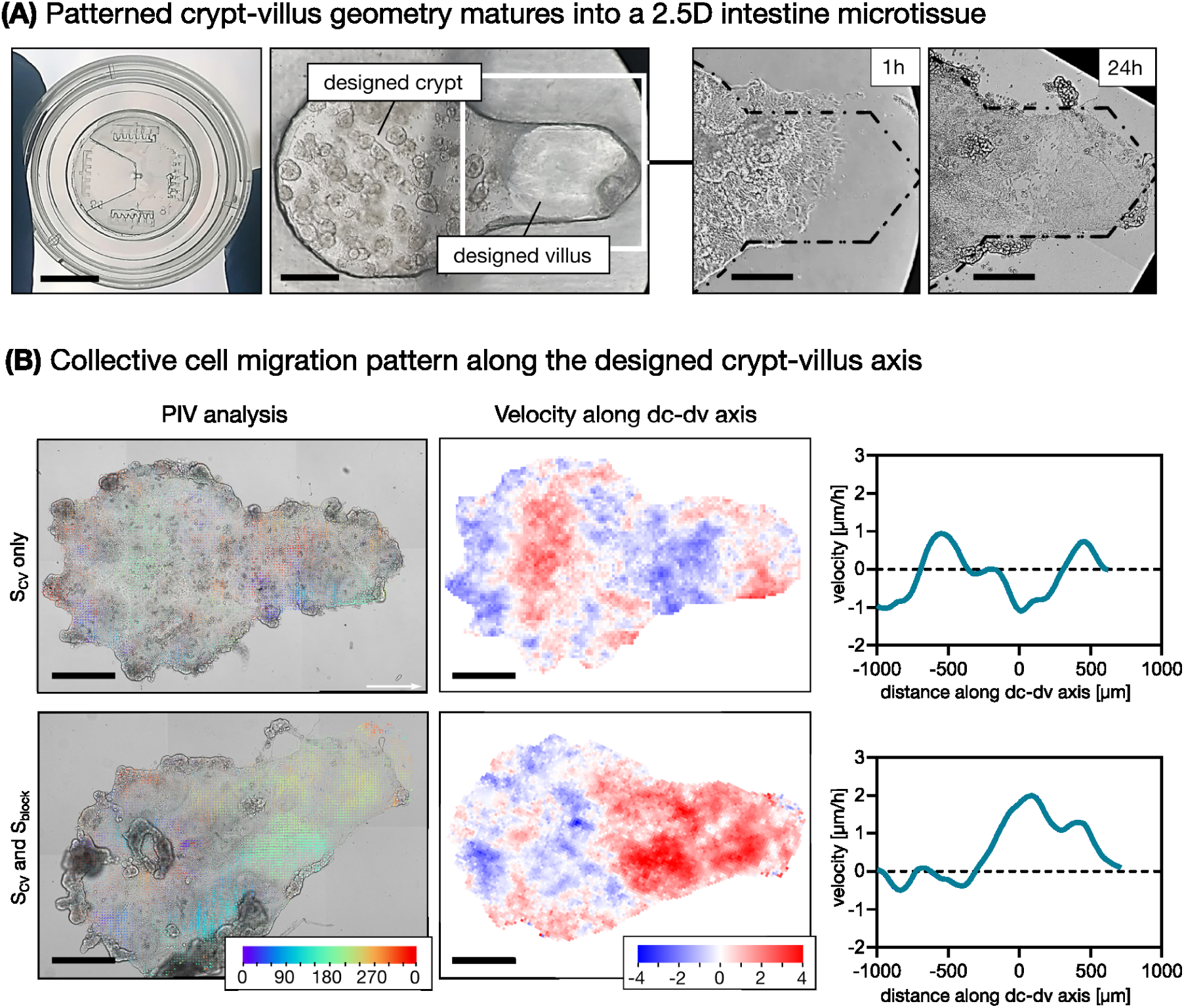
Collective cell migration in patterned intestinal organoid microtissues. **(A)** Both stencils were placed and aligned on the surface of a polyacrylamide gel coated glass bottom dish suited for live cell imaging (left picture, scale bar 5 mm). Intestinal organoids were seeded and cultivated within the crypt zone while S_block_ sealed the villus zone (second brightfield image from the left, scale bar 100 *µ*m). Upon removal of both stencils, cells from the intestinal organoids started to dissociate and migrate towards the designed villus region until full confluency was reached within this region (brightfield images on the right, scale bar 75 µm). **(B)** For particle image velocimetry analysis, mean migration vectors were plotted for 12 hours against the first time point of imaging (scale bar 500 *µ*m). Analysis for directed cell migration revealed a collective migration pattern along the designed crypt-villus axis after full confluency within the villus was reached. In contrast, collective migration was not observed in control samples where organoids were seeded in the whole designated crypt-villus region of stencil S_CV_ without S_block_ obstructing the designed villus region.

## 3 Discussion

We leveraged 3D printing and replica molding for a cost-effective and versatile 2D micropatterning of spatially defined multicellular systems with sub-millimeter resolution. We implemented three applications that showcase various underlying biomedical research questions, each requiring a different spatial cellular arrangement. First, a model of the tumor microenvironment comprising cancer cells and surrounding CAFs was tailored to investigate CAF-mediated resistance to a specific targeted therapy. Second, synthetic morphogen gradients that evolve over space and time were generated in different geometric arrangements. Third, a sequential stenciling approach was used to differentially pattern ECM and primary organoids to model intestinal homeostasis.

Constructing micro-physiological models of native tissues *in vitro* requires a trade-off between the technical complexity inherent to its fabrication and the achievable physiological relevance. A reductionist approach to model tissue structure risks losing essential function, whereas excessive model complexity impedes its accessibility [85]. The cell patterning approach presented here prioritizes cost-effectiveness, ease of use and scalability. Replica-molded PDMS stencils can reliably direct multicellular structures and microtissues in 2D, and can be readily implemented in a life sciences laboratory without the need for complex microfabrication infrastructure. Resulting co-cultures are ideal for high-resolution and live cell imaging. The degree of complexity required from a synthetic microtissue model is directly related to the underlying biological question. In case of the crypt-villus axis, for example, when considering the length scales of an individual intestinal epithelial cell (10 µm) and the average circumference of a villus (300 µm), it becomes apparent that a planar 2D model is a suitable approximation of the tissue geometry. Considering the convex curvature of the villus, a planar model is also a better approximation than a 3D organoid, where the curvature of the central villus-like region is concave. Thus, for addressing questions related to cell migration dynamics in the villus, the crypt-villus microtissue presented here is an appropriate model and offers a far superior platform for high-resolution live imaging compared to 3D organoids.

PDMS elastomers readily absorb non-polar small molecule substances, which is a common concern in many organ-on-chip applications for testing of pharmaceuticals [75]. Unlike patterning strategies that utilize permanent scaffolding [37], PDMS stencils can easily be removed after cell loading and initial cultivation. This minimizes possible interferences with downstream treatment assays. We take advantage of this in our simplified tumor microenvironment design, where a compressive surrounding layer of CAF cells confers resistance of tumor cells to cetuximab, but not afatinib (Figure 4). Previous work suggests that CAF-induced resistance to cetuximab is mediated by increased secretion of HER ligands by CAFs in response to cetuximab treatment [58]. Additional resistance may be conferred by the mechanical compression of the cancer cells by the surrounding CAF layer, which has been shown to induce activation of Yes-associated protein 1 (YAP1) and mechanotransduction [55]. In future studies, this platform can be used to determine how CAFs modulate therapeutic drug responses. This will shed further light on the molecular mechanisms underlying CAF-mediated resistance and can inform treatment strategies based on the stromal reaction. While stencil fabrication itself is efficient, stencil handling and cell loading was performed manually and was hence time-intensive. Future automation, such as robotic stencil placement under sterile conditions and integrated live-cell imaging [90], could achieve high-throughput screens to test multiple cell types and treatment combinations against specific geometric cues.

To demonstrate the potential applications of our microstencilling approach for synthetic morphogenesis, we used two different geometric arrangements of the SynNotch system to generate interfacial morphogen signaling gradients. The rectangular stencil arrangement mimicked previous work inducing gradient formation using commercially available stenciling systems [35]. The concentric circle arrangement illustrates how customized shapes can help investigate the dynamics of organizer centers with different geometries. This can be applied to study pattern formation to test the influence of tissue geometry at distinct morphogenetic events. While this was also previously possible using traditional stereolithography approaches, the incorporation of 3D printing dramatically reduces the time needed to generate new stencil shapes and allows for rapid prototyping. Future studies leveraging this approach may investigate more complex geometries and incorporate more advanced genetic circuits involving multiple cellular components.

Our simplified crypt-villus axis demonstrated that differential patterning of ECM and organoids could be used to impose specific geometries and directional cell migration to achieve homeostatic dynamics that align with previous *in vitro* and *ex vivo* findings [45] (Figure 8). In contrast to previous studies investigating collective cell migration into empty space using micropatterning approaches [46, 47], we analyzed migration dynamics at full confluence. This demonstrated that we were able to achieve a quasi-homeostatic system of cell turnover and migration, similar to the intestinal tissue. Our micropatterned *in vitro* system offers a number of advantages, including customizable geometries and a dramatic reduction in animal use for studies of intestinal cell dynamics. Small intestinal organoids can be grown from tissue harvested from excess breeding mice or from residual tissue from other animal experiments. As organoids from a single animal can be maintained in culture for several months, the relative burden for experimental animals is therefore dramatically reduced. At the same time, the planar micropatterns offer a format which enables high-resolution imaging, making the system ideal for mechanistic studies. Such approaches will prove crucial for further determination of the mechanisms underlying passive and active mechanisms of collective epithelial migration [45]. In addition to our analysis using PIV from brightfield timelapse images, incorporation of organoid lines expressing fluorescent markers will allow for more detailed studies of subcellular dynamics and cell-cell interactions during collective migration.

The flexibility of the synthetic crypt-villus axis will also allow for future investigations of disease mechanisms in which collective motility and tissue geometry are known to be perturbed. Infection of parasitic helminths, for example, is known to cause a significant shortening or lengthening of intestinal villi in distinct regions of the intestine [100]. Chronic intestinal inflammation and the associated increased in inflammatory cytokine production is also known to lead to villus atrophy, resulting in decreased villus length and perturbed migration dynamics [101, 102, 103]. How such changes in tissue geometry influence homeostasis and collective cell migration has thus far not been addressed, likely due to a lack of suitable models. Our crypt-villus axis model presents an ideal platform to understand the relationship between tissue geometry and collective cell dynamics. In addition to using this patterning approach to study cell-level mechanisms, this system can also be upscaled for screening applications, for example to determine how specific small molecules or intestinal microbiota metabolites influence intestinal homeostasis. In future studies, patient-derived organoids can also readily be incorporated into the crypt-villus patterns for fundamental and translational studies of human intestinal pathophysiology.

One of the limitations of 3D printing-based replica molding is the spatial resolution. Higher-resolution techniques as micro-stereolithography [86] or two-photon 3D printing [87] could be adopted to realize finer structures, if needed. For instance, a dual molding process achieved precise PDMS stencil features below 50 µm from precision 2-photon 3D-printed masters [88]. However, stencil adhesion strength is critical to prevent leakage during patterning and subsequent cultivation. This limits how small a cell patch can be reliably patterned. In addition to incorporating other additive manufacturing technologies, the materials used for printing can also be further optimized in the future. For example, parylene-C, thermoplastics or polystyrene films are promising material candidates to evaluate for patterning higher resolution features [89].

## 4 Conclusion

Taken together, our sequential stenciling approach offers a convenient, cost-effective and adaptable workflow for generating specific multicellular geometries. As demonstrated through three selected applications, the flexibility and rapid prototyping enable the generation of organotypic microtissues that can be used to address various biological questions without relying on extensive microfabrication infrastructure. This not only impacts fundamental research, but can also be applied for translational studies and screening applications. As the United States Food and Drug Administration (US FDA) lifted the requirements in 2022 for preclinical in-animal testing, the development of physiologically accurate microtissue models will be central in replacing pre-clinical animal models. The adoption of the sequential stenciling platform may thus prove useful in other biomedical research settings as well.

## 5 Materials and Methods

Unless stated otherwise, all cell culture reagents and culture media were purchased from Thermo Fisher Scientific.

### 5.1 3D-printed master molds

Master molds were designed in SolidWorks (SP 3.0, Dassault Systems) and exported to STL format. Masters were printed in Clear Resin V4 on a Form3 SLA printer with standard settings and 50 µm slicing height configured in Preform software (v3.52.1). Printed molds were developed by washing in isopropanol for 15 minutes using a FormWash and subsequently dried and post-exposure cured for 15 minutes at 37 °C in a FormCure (all Formlabs Inc).

### 5.2 USAF resolution target

3D USAF targets were 3D-printed the same way as master molds, but in addition stained with 1 mg/ml rhodamine B in 50 vol% ethanol in ddH_2_O and for 60 minutes. Targets were imaged at 1200 nm excitation on a Stellaris8-DIVE multiphoton microscope equipped with a 25x NA 0.95 water immersion objective (both Leica microsystems) and an InSightX3 dual laser (Spectra Physics). Brightfield imaging of the whole USAF target was performed using the Axio Imager 2 upright microscope (Zeiss).

### 5.3 PDMS stencil fabrication

Stencils were replica molded in sylgard 184 silicone elastomer (Farnell GmbH). For this, curing agent and base were mixed in 1:20 ratio at 2,000 rpm for 45 seconds, and spun down at 1,500 rpm for 4 minutes uing a planetary ThinkyMixer (Thinky U.S.A). Mixed PDMS was filled into the master molds with a 1 ml syringe and then desiccated for 20 minutes to remove air bubbles trapped in the master molds. PDMS curing proceeded then at 80 °C for 45-50 minutes to achieve the desired curing stage. PDMS stencils were peeled off the master molds and washed multiple times with 100% ethanol.

### 5.4 Cell culturing and stencil loading

All cells were cultivated in a standard cell culture incubator at 37 °C with 5% CO_2_. Prior to cell seeding, PDMS stencils were sterilized with UV light for 60 minutes and placed top-down in a cell culture wellplate or on a collagen-coated coverslip. Next, stencils were filled with cell culture media and preheated at 37° C for at least 60 minutes to prevent gas bubbles formation when adding culture media. Colorectal cancer (LIM1215-mScarlet) and cancer-associated fibroblast (IKP-CAF-001-GFP) cells were cultivated in RPMI-1640 culture medium supplemented with 10% FCS, 1% penicillin/streptomycin. LIM1215-mScarlet culture media was supplemented with 20 µg/ml puromycin (Sigma-Aldrich), IKP-CAF-001-GFP culture media was additionally supplemented with 1% MEM NEAA and 1% GlutaMAX. LIM1215 cells expressing nuclear mScarlet were generated by lentiviral transduction with pLV-EF1*α*-NLS-mScarlet-IRES-Puro, followed by puromycin selection. For circular LIM1215 cell patches, 1.000 LIM1215-mScarlet cells in 2 µl media volume were seeded in the stencil opening. For CRC-CAF co-cultures 20 µl with about 17.000 IKP-CAF-001-GFP or LIM1215-GFP control cells were seeded in the outer stencil. Cancer cells were incubated for 3 hours for an initial cell attachment before removing the inner stencil. Both CRC and CAF cells were then let mature for 24 hours. For co-cultures, IKP-CAF-001-GFP culture media was used.

The construction of GFP-secreting “sender cells” and GFP-receiving “receiver cells” is described in [91]. Briefly, mouse L929 fibroblasts (RIKEN BRC, #RCB1422) were lenti-virally transduced to create both the sender and receiver cells. The sender cells express a secretory GFP that is a fusion protein of the Gaussialuciferase signal sequence and EGFP. The receiver cells express anti-GFP synNotch receptor (Addgene, #162230) and GFP-anchor protein (Addgene, #162225) to sense GFP and induce the expression of mCherry fluorescence reporter. These cells were maintained in DMEM culture medium supplemented with 10% FCS and 1% penicillin/streptomycin.

Generation of organoids was performed as previously described [92]. Briefly, organoids were isolated from the intestines of C57BL/6 mice younger then 6-month-old by resecting the small intestine from the stomach to the caecum, flushing its contents, cutting longitudinally, and washing on ice in PBS^−/−^ with 2x Antibiotic/Antimycotic (Anti/Anti). The tissue was minced, incubated for 30 minutes in 2 mM EDTA in PBS^−/−^ with 2x Anti/Anti at 4 °C on a rocking shaker and resuspended in PBS^−/−^ with 2x Anti/Anti. All plasticware was coated with 10% FCS in PBS^−/−^ to avoid losing crypts due to non-specific adhesion to plastic. Tissue pieces were sequentially pipetted to liberate the crypts from the epithelial tissue. The supernatant of the first 2 fractions was villus-rich and discarded, and subsequent crypt-enriched supernatants were collected. Crypt suspensions were filtered with a 100 *µ*m cell strainer and centrifuged to remove single cells. The pellet was resuspended in DMEM/F-12 with 2x Anti/Anti, filtered with a 70 *µ*M cell strainer, and centrifuged again. The pellet was resuspended in a 1:1 mixture of DMEM/F-12 with 2x Anti/Anti and Basement Membrane Extract Type 2 (BME2) (R&D Systems). A total of 50 µl domes were plated in a 24-well plate, polymerized for 45 minutes at 37 °C and 5% CO_2_. The small intestinal organoids were cultured in DMEM/F-12 medium supplemented with 20 ng/ml mEGF, 100 ng/ml mFGF, 1x B27, 2x N_2_ with 10% R-spondin and 10% Noggin conditioned medium added (ENR medium). Noggin conditioned medium was produced using the noggin Fc HEK cell line [93] and R-spondin conditioned medium using HA-R-Spondin1-Fc 293T cells (bio-techne). To support stem cell growth, 1 *µ*M Y-27632 (Tebu-Bio) and 3 *µ*M CHIR-99021 (Sigma-Aldrich) were added to the ENR medium. After one day, the medium was changed to standard ENR medium and crypts developed into organoids for five days before the first passage. Organoids were cultured in 3D before plating into 2D organoid monolayers for migration analysis.

### 5.5 Immunofluorescence imaging

For immunofluorescence staining, CRC-CAF and CRC-CRC co-cultures seeded on collagen-coated coverslips were incubated in a standard culture incubator at 5% CO_2_ at 37 °C for the corresponding time. Next, cells were washed with 1x PBS, fixed with 4% paraformaldehyde (PFA) for 15 minutes, and afterwards washed three times with 1x PBS for 15 minutes. After washing, cells were permeabilized with 0.1% Triton X-100 in 1x PBS for 5 minutes. To block non-specific binding, coverslips were incubated with 5% FCS in 1x PBS for 30 minutes at room temperature and subsequently incubated with 4,6-diamidino-2-phenylindole (DAPI) and Alexa Fluor 633-Phalloidin, dissolved in 1x PBS and 5% FCS for 1 hour. For live-dead staining, cells were incubated in RPMI1640 medium without serum and 1 µM Calcein/AM for 30 minutes and fixed afterwards. Immunofluorescence imaging of fixed and stained cell samples was performed with a Stellaris Dive 8 multiphoton microscope (Leica) using either a 10x, NA 0.4 or 25x, NA 0.95 water-immersion objective. DAPI, mScarlet, GFP, rhodamine B and AlexaFluor633-Phalloidin were excited at 1045 nm, 1150 nm, 900 nm, 1200 nm and 1250 nm, respectively.

### 5.6 Anti-tumor drug treatment and analysis

CRC-CAF and CRC-CRC co-cultures were either seeded in 96-well plates or on collagen-coated coverslips. For this, coverslips were incubated with a 1:200 collagen solution in 1x PBS at 37 °C for 120 minutes beforehand. Co-cultures were cultivated until full confluency was reached. Next, the culture medium was discarded and exchanged for 200 µl of fresh culture medium with the drug. For targeting the LIM1215-mScarlet cancer cells, 100 nM cetuximab, dissolved in 1x PBS, or 100 nM afatinib, dissolved in dimethyl sulfoxide (DMSO), was applied. CRC-CAF and CRC-CRC co-cultures were cultivated for 3 days upon treatment with cetuximab and afatinib. Cultivation was performed inside the IncuCyte S3 (Sartorius) live-cell analysis system at 5% CO_2_ at 37 °C for live-cell imaging of CRC-CAF and CRC-CRC co-cultures that were seeded in 96-well plates. Pictures were captured every 6 hours for the corresponding cultivation time. Obtained images from live-cell imaging and confocal fluorescence imaging were analyzed via FIJI (ImageJ) [94] using the Analyze Particles command thresholded for the lowest common fluorescence value for region of interest (ROI) generation. The relative and absolute areas and the mean mScarlet- as well as GFP-fluorescence intensity signal were quantified in the defined ROI.

### 5.7 synNotch gradient formation

To create a linear gradient pattern within a monolayer of receiver cells, we placed a stencil containing two 4 mm x 8 mm rectangle wells (S_L_, design 5) at the center of wells in a 12-wellplate. About 1.6x10^4^ sender or receiver cells in 32 µl of medium were seeded into the two wells inside the stencil to form adjacent tissues of sender and receiver cells. Then 1.5x10^5^ receiver cells were seeded outside the stencil to serve as a sink. After 4 hours of incubation, the stencil was removed and replaced the media with a mixed media of DMEM/PBS containing 10% FCS and 1% agarose. Once the medium solidified, synNotch signaling was imaged every 2 hr for 48 hr using an Incucyte S3 (Sartorius). To create a radial gradient pattern, a stencil containing a round 2 mm radius circular well (S_R_, design 6) was placed at the center of wells in a 12-well plate. 1.25x10^4^ sender cells were seeded into the circular well of the stencil. After overnight incubation, stencil was removed and 1x10^5^ receiver cells were seeded into the well. After 4 hours of cultivation, the medium was replaced with the DMEM/PBS containing 10% FCS and 1% agarose and imaged every 2 hours for a total of 50 hours using an Incucyte S3.

### 5.8 Live cell imaging and cell migration analysis within intestinal organoid microtissues

For generating organoid microtissues, 35 mm glass bottom dishes were pretreated with Silane APTMS diluted 1:2 in ddH_2_O for 15 minutes, washed with ddH_2_O, and incubated with 0.5% Glutaraldehyde in PBS for 30 minutes. The glass bottom dishes were then briefly washed with ddH_2_O and dried. 5 kPa polyacrylamide gels were polymerized on the 35 mm glass bottom dishes and functionalized with 3-4-Dihydroxy-L-phenylalanine (L-Dopa, Sigma-Aldrich). The functionalized polyacrylamide gels were coated with 250 *µ*g/ml Rat Tail Collagen-I (Corning) and 250 *µ*g/ml Laminin 1 (Sigma-Aldrich) in 10 mM HEPES using the S_CV_ stencil to pattern Rat Tail Collagen-I and Laminin 1 deposition into the crypt-villus axis desiccating for 1 minute to remove air bubbles and incubated overnight at 4 °C. One day before plating, organoids were passaged and on the day of plating organoids were freed from BME2 by pipetting and incubating in harvesting solution (bio-techne) for 1 hour at 4 °C. Freed crypts were resuspended in ENR medium and S_block_ was aligned on S_CV_. Intestinal crypts were seeded into the crypt zone of S_CV_ and incubated for 1 hour at 37 °C and 5% CO_2_ for initial attachment. Both stencils were carefully peeled off the polyacrylamide gel surface to open the villus zone for cell migration, and 1 ml of ENR medium with 2x Anti/Anti was overlaid. Organoid microtissues were cultivated until they reached full confluency in the crypt-villus axis before migration analysis.

Organoid monolayers were supplemented with 2 ml of ENR + 10% FCS + 1% Glutamax + ciprofloxacin (5 µg/ml, Sigma-Aldrich) + metronidazole (1 µg/ml, Sigma-Aldrich) and imaged at full confluency using the Axio Cell Observer (Zeiss). Images were captured in 30 min intervals for 12 hours. Cell migration analysis of differentiating cells was performed with particle image velocimetry [95] using the following packages: numpy [96], openpiv [99] and matplotlib [97].

## 6 Acknowledgements

The authors gratefully acknowledge the Technology Platform “Cellular Analytics” of the Stuttgart Research Center Systems Biology for their support and assistance in this work. We thank Stephan Eisler and Melanie Noack for their technical assistance with multiphoton and live-cell imaging. We also thank the staff of the animal facility of the Institute of Cell Biology and Immunology at the University of Stuttgart for their care of mouse stocks and support for experimental procedures. We thank Karen Kresbach for help with establishing intestinal organoid cell culture and first stencil designs during early stages of the project. The pLV-EF1*α*-NLS-mScarlet-IRES-Puro plasmid and lentiviral particles were kindly provided by Dr. Cristiana Lungu, University of Stuttgart. Noggin Fc HEK cells were kindly provided by Jacqueline Vermeulen and Vanesa Muncan, University Medical Clinic Amsterdam. IKP-CAF-001-GFP cells were kindly provided by Dr. Meng Dong, Dr. Margarete Fischer-Bosch Institute of Clinical Pharmacology.

This work was supported by the Ministry of Science, Research and Arts Baden-Württemberg within the 3R-US project (33-7533-6-1522/2/3). J.C. and A.G.C. are supported by the Federal Ministry of Education and Research (BMBF) and the Baden-Württemberg Ministry of Science (MWK-BW) as part of the Excellence Strategy of the German Federal and State Governments (NWG-GastroTumors to A.G.C.) S.T. was supported by the Japan Science and Technology Agency (JST), PRESTO Grant No. JPMJPR2147 and FOREST Grant No. JPMJFR2311 and the Japan Society for the Promotion of Science (JSPS), KAKENHI Grant No.21H05291 and 24K0202.

## Notes

### Competing Interest Statement

The authors have declared no competing interest.

## References

[1] Zhu M, Zernicka-Goetz M. Principles of Self-Organization of the Mammalian Embryo. Cell. 2020 Dec 10;183(6):1467–1478. doi: 10.1016/j.cell.2020.11.003.

[2] Schnell S, Painter K J, Maini P K and Othmer H G 2001 Spatiotemporal pattern formation in early development: a review of primitive streak formation and somitogenesis Mathematical Models for Biological Pattern Formation pp 11–37. doi: 10.1007/978-1-4613-0133-2_2

[3] Guller A, Igrunkova A. Engineered Microenvironments for 3D Cell Culture and Regenerative Medicine: Challenges, Advances, and Trends. Bioengineering (Basel). 2022 Dec 22;10(1):17. doi: 10.3390/bioengineering10010017.

[4] Masaoutis C, Theocharis S. The Role of Exosomes in Bone Remodeling: Implications for Bone Physiology and Disease. Dis Markers. 2019 Aug 14;2019:9417914. doi: 10.1155/2019/9417914.

[5] Moro LG, Guarnier LP, Azevedo MF, Fracasso JAR, Lucio MA, Castro MV, Dias ML, Lívero FADR, Ribeiro-Paes JT. A Brief History of Cell Culture: From Harrison to Organs-on-a-Chip. Cells. 2024 Dec 15;13(24):2068. doi: 10.3390/cells13242068.

[6] Duval K, Grover H, Han LH, Mou Y, Pegoraro AF, Fredberg J, Chen Z. Modeling Physiological Events in 2D vs. 3D Cell Culture. Physiology (Bethesda). 2017 Jul;32(4):266–277. doi: 10.1152/physiol.00036.2016.

[7] Jensen C, Teng Y. Is It Time to Start Transitioning From 2D to 3D Cell Culture? Front Mol Biosci. 2020 Mar 6;7:33. doi: 10.3389/fmolb.2020.00033.

[8] Yamaguchi S, Kaneko M, Narukawa M. Approval success rates of drug candidates based on target, action, modality, application, and their combinations. Clin Transl Sci. 2021 May;14(3):1113–1122. doi: 10.1111/cts.12980. Epub 2021 Apr 8.

[9] Cordeiro S, Oliveira BB, Valente R, Ferreira D, Luz A, Baptista PV, Fernandes AR. Breaking the mold: 3D cell cultures reshaping the future of cancer research. Front Cell Dev Biol. 2024 Nov 26;12:1507388. doi: 10.3389/fcell.2024.1507388.

[10] Pinto B, Henriques AC, Silva PMA, Bousbaa H. Three-Dimensional Spheroids as In Vitro Pre-clinical Models for Cancer Research. Pharmaceutics. 2020 Dec 6;12(12):1186. doi: 10.3390/pharmaceutics12121186.

[11] Wang Y, Gao Y, Pan Y, Zhou D, Liu Y, Yin Y, Yang J, Wang Y, Song Y. Emerging trends in organ-on-a-chip systems for drug screening. Acta Pharm Sin B. 2023 Jun;13(6):2483–2509. doi: 10.1016/j.apsb.2023.02.006. Epub 2023 Feb 15.

[12] Xiang Y, Miller K, Guan J, Kiratitanaporn W, Tang M, Chen S. 3D bioprinting of complex tissues in vitro: state-of-the-art and future perspectives. Arch Toxicol. 2022 Mar;96(3):691–710. doi: 10.1007/s00204-021-03212-y. Epub 2022 Jan 10.

[13] Goubko CA, Cao X. Patterning multiple cell types in co-cultures: A review. Mater Sci Eng C Mater Biol Appl. 2009 Aug;29(6):1855–1868. doi: 10.1016/j.msec.2009.02.016

[14] Guguen-Guillouzo C, Clèment B, Baffet G, Beaumont C, Morel-Chany E, Glaise D, Guillouzo A. Maintenance and reversibility of active albumin secretion by adult rat hepatocytes co-cultured with another liver epithelial cell type. Exp Cell Res. 1983 Jan;143(1):47–54. doi: 10.1016/0014-4827(83)90107-6.

[15] Shimaoka S, Nakamura T, Ichihara A. Stimulation of growth of primary cultured adult rat hepatocytes without growth factors by coculture with nonparenchymal liver cells. Exp Cell Res. 1987 Sep;172(1):228–42. doi: 10.1016/0014-4827(87)90109-1.

[16] Schrode W, Mecke D, Gebhardt R. Induction of glutamine synthetase in periportal hepatocytes by cocultivation with a liver epithelial cell line. Eur J Cell Biol. 1990 Oct;53(1):35–41.

[17] Chiu DT, Jeon NL, Huang S, Kane RS, Wargo CJ, Choi IS, Ingber DE, Whitesides GM. Patterned deposition of cells and proteins onto surfaces by using three-dimensional microfluidic systems. Proc Natl Acad Sci U S A. 2000 Mar 14;97(6):2408–13. doi: 10.1073/pnas.040562297.

[18] Tien J, Nelson CM, Chen CS. Fabrication of aligned microstructures with a single elastomeric stamp. Proc Natl Acad Sci U S A. 2002 Feb 19;99(4):1758–62. doi: 10.1073/pnas.042493399. Epub 2002 Feb 12.

[19] Bhatia SN, Balis UJ, Yarmush ML, Toner M. Probing heterotypic cell interactions: hepatocyte function in microfabricated co-cultures. J Biomater Sci Polym Ed. 1998;9(11):1137–60. doi: 10.1163/156856298x00695.

[20] Hui EE, Bhatia SN. Micromechanical control of cell-cell interactions. Proc Natl Acad Sci U S A. 2007 Apr 3;104(14):5722–6. doi: 10.1073/pnas.0608660104. Epub 2007 Mar 27.

[21] Takahashi S, Yamazoe H, Sassa F, Suzuki H, Fukuda J. Preparation of coculture system with three extracellular matrices using capillary force lithography and layer-by-layer deposition. J Biosci Bioeng. 2009 Dec;108(6):544–50. doi: 10.1016/j.jbiosc.2009.06.013. Epub 2009 Jul 24.

[22] Yamazoe H, Ichikawa T, Hagihara Y, Iwasaki Y. Generation of a patterned co-culture system composed of adherent cells and immobilized nonadherent cells. Acta Biomater. 2016 Feb;31:231–240. doi: 10.1016/j.actbio.2015.12.016. Epub 2015 Dec 11.

[23] Ayibaike D, Cui M, Wei J. Single-cell patterning based on immunocapture and a surface modified substrate. Appl Sci. 2018;8:2152. doi: 10.3390/app8112152

[24] Fukuda J, Khademhosseini A, Yeh J, Eng G, Cheng J, Farokhzad OC, Langer R. Micropatterned cell co-cultures using layer-by-layer deposition of extracellular matrix components. Biomaterials. 2006 Mar;27(8):1479–86. doi: 10.1016/j.biomaterials.2005.09.015. Epub 2005 Oct 19.

[25] Kim SM, Fukuda J, Khademhosseini A. Patterned cocultures for controlling cell-cell interactions. In: Khademhosseini A, Borenstein J, Toner M, Takayama S, editors. Micro- and Nanoengineering of the Cellular Microenvironment. Norwood (MA): Artech House; 2006. p. 53–63

[26] Kaji H, Yokoi T, Kawashima T, Nishizawa M. Controlled cocultures of HeLa cells and human umbilical vein endothelial cells on detachable substrates. Lab Chip. 2009 Feb 7;9(3):427–32. doi: 10.1039/b812510d. Epub 2008 Nov 6.

[27] Wright D, Rajalingam B, Selvarasah S, Dokmeci MR, Khademhosseini A. Generation of static and dynamic patterned co-cultures using microfabricated parylene-C stencils. Lab Chip. 2007 Oct;7(10):1272–9. doi: 10.1039/b706081e. Epub 2007 Jul 25.

[28] Hassanzadeh-Barforoushi A, Shemesh J, Farbehi N, Asadnia M, Yeoh GH, Harvey RP, Nordon RE, Warkiani ME. A rapid co-culture stamping device for studying intercellular communication. Sci Rep. 2016 Oct 18;6:35618. doi: 10.1038/srep35618.

[29] Gimenez R, Pérez-Sosa C, Bourguignon N, Miriuka S, Bhansali S, Arroyo CR, Debut A, Lerner B, Pérez MS. Simple Microcontact Printing Technique to Obtain Cell Patterns by Lithography Using Grayscale, Photopolymer Flexographic Mold, and PDMS. Biomimetics (Basel). 2022 Oct 8;7(4):155. doi: 10.3390/biomimetics7040155.

[30] Chiu DT, Jeon NL, Huang S, Kane RS, Wargo CJ, Choi IS, Ingber DE, Whitesides GM. Patterned deposition of cells and proteins onto surfaces by using three-dimensional microfluidic systems. Proc Natl Acad Sci U S A. 2000 Mar 14;97(6):2408–13. doi: 10.1073/pnas.040562297.

[31] Kicheva A, Briscoe J. Control of Tissue Development by Morphogens. Annu Rev Cell Dev Biol. 2023 Oct 16;39:91–121. doi: 10.1146/annurev-cellbio-020823-011522. Epub 2023 Jul 7.

[32] Pinheiro D, Kardos R, Hannezo E, Heisenberg CP. Morphogen gradient orchestrates pattern-preserving tissue morphogenesis via motility-driven unjamming. Nat Phys. 2022;18:1482–1493. doi: 10.1038/s41567-022-01767-8 no pubmed

[33] Simsek MF, Özbudak EM. Patterning principles of morphogen gradients. Open Biol. 2022 Oct;12(10):220224. doi: 10.1098/rsob.220224. Epub 2022 Oct 19.

[34] Morsut L, Roybal KT, Xiong X, Gordley RM, Coyle SM, Thomson M, Lim WA. Engineering Customized Cell Sensing and Response Behaviors Using Synthetic Notch Receptors. Cell. 2016 Feb 11;164(4):780–91. doi: 10.1016/j.cell.2016.01.012. Epub 2016 Jan 28.

[35] Toda S, Blauch LR, Tang SKY, Morsut L, Lim WA. Programming self-organizing multicellular structures with synthetic cell-cell signaling. Science. 2018 Jul 13;361(6398):156–162. doi: 10.1126/science.aat0271. Epub 2018 May 31.

[36] Toda S, McKeithan WL, Hakkinen TJ, Lopez P, Klein OD, Lim WA. Engineering synthetic morphogen systems that can program multicellular patterning. Science. 2020 Oct 16;370(6514):327–331. doi: 10.1126/science.abc0033.

[37] Garibyan M, Hoffman T, Makaske T, Do SK, Wu Y, Williams BA, March AR, Cho N, Pedroncelli N, Lima RE, Soto J, Jackson B, Santoso JW, Khademhosseini A, Thomson M, Li S, McCain ML, Morsut L. Engineering programmable material-to-cell pathways via synthetic notch receptors to spatially control differentiation in multicellular constructs. Nat Commun. 2024 Jul 13;15(1):5891. doi: 10.1038/s41467-024-50126-1. Erratum in: Nat Commun. 2024 Aug 1;15(1):6490. doi: 10.1038/s41467-024-50845-5.

[38] Kwon O, Han TS, Son MY. Intestinal Morphogenesis in Development, Regeneration, and Disease: The Potential Utility of Intestinal Organoids for Studying Compartmentalization of the Crypt-Villus Structure. Front Cell Dev Biol. 2020 Oct 23;8:593969. doi: 10.3389/fcell.2020.593969.

[39] Umar S. Intestinal stem cells. Curr Gastroenterol Rep. 2010 Oct;12(5):340–8. doi: 10.1007/s11894-010-0130-3.

[40] De Gregorio V, Imparato G, Urciuolo F, Netti PA. Micro-patterned endogenous stroma equivalent induces polarized crypt-villus architecture of human small intestinal epithelium. Acta Biomater. 2018 Nov;81:43–59. doi: 10.1016/j.actbio.2018.09.061. Epub 2018 Sep 30.

[41] Altay G, Larrañaga E, Tosi S, Barriga FM, Batlle E, Fernández-Majada V, Martínez E. Self-organized intestinal epithelial monolayers in crypt and villus-like domains show effective barrier function. Sci Rep. 2019 Jul 12;9(1):10140. doi: 10.1038/s41598-019-46497-x. Erratum in: Sci Rep. 2019 Dec 6;9(1):18822. doi: 10.1038/s41598-019-55181-z.

[42] Tian CM, Yang MF, Xu HM, Zhu MZ, Yue NN, Zhang Y, Shi RY, Yao J, Wang LS, Liang YJ, Li DF. Stem cell-derived intestinal organoids: a novel modality for IBD. Cell Death Discov. 2023 Jul 21;9(1):255. doi: 10.1038/s41420-023-01556-1.

[43] Shin W, Wu A, Min S, Shin YC, Fleming RYD, Eckhardt SG, Kim HJ. Spatiotemporal Gradient and Instability of Wnt Induce Heterogeneous Growth and Differentiation of Human Intestinal Organoids. iScience. 2020 Aug 21;23(8):101372. doi: 10.1016/j.isci.2020.101372. Epub 2020 Jul 16.

[44] Heymann M, Fraden S, Kim D. Multi-height precision alignment with selectively developed alignment marks. J Microelectromech Syst. 2014;23:424–427. doi: 10.1109/JMEMS.2013.2290974

[45] Krndija D, El Marjou F, Guirao B, Richon S, Leroy O, Bellaiche Y, Hannezo E, Matic Vignjevic D. Active cell migration is critical for steady-state epithelial turnover in the gut. Science. 2019 Aug 16;365(6454):705–710. doi: 10.1126/science.aau3429.

[46] Vedula SR, Leong MC, Lai TL, Hersen P, Kabla AJ, Lim CT, Ladoux B. Emerging modes of collective cell migration induced by geometrical constraints. Proc Natl Acad Sci U S A. 2012 Aug 7;109(32):12974–9. doi: 10.1073/pnas.1119313109. Epub 2012 Jul 19.

[47] Tarle V, Gauquelin E, Vedula SRK, D’Alessandro J, Lim CT, Ladoux B, Gov NS. Modeling collective cell migration in geometric confinement. Phys Biol. 2017 May 3;14(3):035001. doi: 10.1088/1478-3975/aa6591.

[48] Tonini L, Ahn C. Latest Advanced Techniques for Improving Intestinal Organoids Limitations. Stem Cell Rev Rep. 2025 Aug;21(6):1631–1647. doi: 10.1007/s12015-025-10894-9. Epub 2025 May 19.

[49] Mitrofanova O, Nikolaev M, Xu Q, Broguiere N, Cubela I, Camp JG, Bscheider M, Lutolf MP. Bioengineered human colon organoids with in vivo-like cellular complexity and function. Cell Stem Cell. 2024 Aug 1;31(8):1175–1186.e7. doi: 10.1016/j.stem.2024.05.007. Epub 2024 Jun 13.

[50] Verhulsel M, Simon A, Bernheim-Dennery M, Gannavarapu VR, Gérémie L, Ferraro D, Krndija D, Talini L, Viovy JL, Vignjevic DM, Descroix S. Developing an advanced gut on chip model enabling the study of epithelial cell/fibroblast interactions. Lab Chip. 2021 Jan 21;21(2):365–377. doi: 10.1039/d0lc00672f. Epub 2020 Dec 11. Erratum in: Lab Chip. 2023 Mar 14;23(6):1713. doi: 10.1039/d3lc90020g.

[51] Nikolaev M, Mitrofanova O, Broguiere N, Geraldo S, Dutta D, Tabata Y, Elci B, Brandenberg N, Kolotuev I, Gjorevski N, Clevers H, Lutolf MP. Homeostatic mini-intestines through scaffold-guided organoid morphogenesis. Nature. 2020 Sep;585(7826):574–578. doi: 10.1038/s41586-020-2724-8. Epub 2020 Sep 16.

[52] Zafari N, Khosravi F, Rezaee Z, Esfandyari S, Bahiraei M, Bahramy A, Ferns GA, Avan A. The role of the tumor microenvironment in colorectal cancer and the potential therapeutic approaches. J Clin Lab Anal. 2022 Aug;36(8):e24585. doi: 10.1002/jcla.24585. Epub 2022 Jul 8.

[53] Mayer S, Milo T, Isaacson A, Halperin C, Miyara S, Stein Y, Lior C, Pevsner-Fischer M, Tzahor E, Mayo A, Alon U, Scherz-Shouval R. The tumor microenvironment shows a hierarchy of cell-cell interactions dominated by fibroblasts. Nat Commun. 2023 Sep 19;14(1):5810. doi: 10.1038/s41467-023-41518-w.

[54] Tao L, Huang G, Song H, Chen Y, Chen L. Cancer associated fibroblasts: An essential role in the tumor microenvironment. Oncol Lett. 2017 Sep;14(3):2611–2620. doi: 10.3892/ol.2017.6497. Epub 2017 Jun 30.

[55] Barbazan J, Pérez-González C, Gómez-González M, Dedenon M, Richon S, Latorre E, Serra M, Mariani P, Descroix S, Sens P, Trepat X, Vignjevic DM. Cancer-associated fibroblasts actively compress cancer cells and modulate mechanotransduction. Nat Commun. 2023 Nov 1;14(1):6966. doi: 10.1038/s41467-023-42382-4.

[56] Feng B, Wu J, Shen B, Jiang F, Feng J. Cancer-associated fibroblasts and resistance to anticancer therapies: status, mechanisms, and countermeasures. Cancer Cell Int. 2022 Apr 29;22(1):166. doi: 10.1186/s12935-022-02599-7.

[57] Wright K, Ly T, Kriet M, Czirok A, Thomas SM. Cancer-Associated Fibroblasts: Master Tumor Microenvironment Modifiers. Cancers (Basel). 2023 Mar 22;15(6):1899. doi: 10.3390/cancers15061899.

[58] Garvey CM, Lau R, Sanchez A, Sun RX, Fong EJ, Doche ME, Chen O, Jusuf A, Lenz HJ, Larson B, Mumenthaler SM. Anti-EGFR Therapy Induces EGF Secretion by Cancer-Associated Fibroblasts to Confer Colorectal Cancer Chemoresistance. Cancers (Basel). 2020 May 28;12(6):1393. doi: 10.3390/cancers12061393.

[59] Feng Y, Ma W, Zang Y, Guo Y, Li Y, Zhang Y, Dong X, Liu Y, Zhan X, Pan Z, Luo M, Wu M, Chen A, Kang D, Chen G, Liu L, Zhou J, Zhang R. Spatially organized tumor-stroma boundary determines the efficacy of immunotherapy in colorectal cancer patients. Nat Commun. 2024 Nov 26;15(1):10259. doi: 10.1038/s41467-024-54710-3.

[60] Souza da Silva R, Queiroga EM, de Toledo Osório C, Cunha KS, Neves FP, Andrade JP, Dias EP. Expression Profile of Microenvironmental Factors in the Interface Zone of Colorectal Cancer: Histological-Stromal Biomarkers and Cancer Cell-Cancer-Associated Fibroblast-Related Proteins Combined for the Assessment of Tumor Progression. Pathobiology. 2024;91(2):99–107. doi: 10.1159/000531695. Epub 2023 Jun 27.

[61] Photronics I, Yomazzo M, Van Den Broeke D. Photomask requirements needed to support future lithography. Microelectron Eng. 1998;41:53–58

[62] LaFratta CN, Baldacchini T, Farrer RA, Fourkas JT, Teich MC, Shih MY, Saleh BEA. Replication of two-photon–polymerized structures for microfabrication. Proc Natl Acad Sci U S A. 2006;103(23):8589–8594. doi: 10.1073/pnas.0602713103

[63] Hagemann C, Bailey MCD, Carraro E, Stankevich KS, Lionello VM, Khokhar N, Suklai P, Moreno-Gonzalez C, O’Toole K, Konstantinou G, Dix CL, Joshi S, Giagnorio E, Bergholt MS, Spicer CD, Imbert A, Tedesco FS, Serio A. Low-cost, versatile, and highly reproducible microfabrication pipeline to generate 3D-printed customised cell culture devices with complex designs. PLoS Biol. 2024 Mar 13;22(3):e3002503. doi: 10.1371/journal.pbio.3002503.

[64] Sun HB, Kawata S. Two-photon photopolymerization and 3D lithographic microfabrication. NMR Biomed. 2004;1:65–85

[65] Bougdid Y, Sekkat Z. Voxels Optimization in 3D Laser Nanoprinting. Sci Rep. 2020 Jun 26;10(1):10409. doi: 10.1038/s41598-020-67184-2.

[66] Bagheri A, Jin J. Photopolymerization in 3D printing. ACS Appl Polym Mater. 2019;1:593–611. doi: 10.1021/acsapm.8b00165

[67] Yen A. Rayleigh or Abbe? Origin and naming of the resolution formula of microlithography. J Micro/Nanolith MEMS MOEMS. 2020;19:040501. doi: 10.1117/1.JMM.19.4.040501

[68] Zhou S, Jiang L. A modern description of Rayleigh’s criterion. Phys Rev A. 2019;99:013808. doi: 10.1103/PhysRevA.99.013808

[69] Stamnes JJ. Focusing of a perfect wave and the Airy pattern formula. Opt Commun. 1981;37:311–314. doi: 10.1016/0030-4018(81)90101-9

[70] Blom H, Driessen J, van den Berg J. The use of a Siemens star to quantify lateral resolution in wide-field fluorescence microscopy. J Microsc. 2008;232:195–204. doi: 10.1111/j.1365-2818.2008.02057.

[71] Sanli UT, Jiao C, Baluktsian M, Grévent C, Hahn K, Wang Y, Srot V, Richter G, Bykova I, Weigand M, Schütz G, Keskinbora K. 3D Nanofabrication of High-Resolution Multilayer Fresnel Zone Plates. Adv Sci (Weinh). 2018 Jun 5;5(9):1800346. doi: 10.1002/advs.201800346.

[72] Broxton M, Grosenick L, Yang S, Cohen N, Andalman A, Deisseroth K, Levoy M. Wave optics theory and 3-D deconvolution for the light field microscope. Opt Express. 2013 Oct 21;21(21):25418–39. doi: 10.1364/OE.21.025418.

[73] Kim GM, Lee SJ, Kim CL. Assessment of the Physical, Mechanical, and Tribological Properties of PDMS Thin Films Based on Different Curing Conditions. Materials (Basel). 2021 Aug 10;14(16):4489. doi: 10.3390/ma14164489.

[74] Venzac B. Light-based 3D printing and post-treatments of moulds for PDMS soft lithography. Lab Chip. 2025 Apr 29;25(9):2129–2147. doi: 10.1039/d4lc00836g.

[75] Toepke MW, Beebe DJ. PDMS absorption of small molecules and consequences in microfluidic applications. Lab Chip. 2006;6:1484–1486. doi: 10.1039/b612140c

[76] Regehr KJ, Domenech M, Koepsel JT, Carver KC, Ellison-Zelski SJ, Murphy WL, Schuler LA, Alarid ET, Beebe DJ. Biological implications of polydimethylsiloxane-based microfluidic cell culture. Lab Chip. 2009 Aug 7;9(15):2132–9. doi: 10.1039/b903043c. Epub 2009 Jun 4.

[77] Xie YH, Chen YX, Fang JY. Comprehensive review of targeted therapy for colorectal cancer. Signal Transduct Target Ther. 2020 Mar 20;5(1):22. doi: 10.1038/s41392-020-0116-z.

[78] Ventola CL. Pharmacogenomics in clinical practice: reality and expectations. P T. 2011 Jul;36(7):412–50.

[79] De Pauw I, Wouters A, Van den Bossche J, Peeters M, Pauwels P, Deschoolmeester V, Vermorken JB, Lardon F. Preclinical and clinical studies on afatinib in monotherapy and in combination regimens: Potential impact in colorectal cancer. Pharmacol Ther. 2016 Oct;166:71–83. doi: 10.1016/j.pharmthera.2016.06.014. Epub 2016 Jun 29.

[80] Assimakopoulos SF, Triantos C, Maroulis I, Gogos C. The Role of the Gut Barrier Function in Health and Disease. Gastroenterology Res. 2018 Aug;11(4):261–263. doi: 10.14740/gr1053w. Epub 2018 Feb 8.

[81] Kasendra M, Tovaglieri A, Sontheimer-Phelps A, Jalili-Firoozinezhad S, Bein A, Chalkiadaki A, Scholl W, Zhang C, Rickner H, Richmond CA, Li H, Breault DT, Ingber DE. Development of a primary human Small Intestine-on-a-Chip using biopsy-derived organoids. Sci Rep. 2018 Feb 13;8(1):2871. doi: 10.1038/s41598-018-21201-7.

[82] Shin W, Kim HJ. 3D in vitro morphogenesis of human intestinal epithelium in a gut-on-a-chip or a hybrid chip with a cell culture insert. Nat Protoc. 2022 Mar;17(3):910–939. doi: 10.1038/s41596-021-00674-3. Epub 2022 Feb 2.

[83] Moorefield EC, Blue RE, Quinney NL, Gentzsch M, Ding S. Generation of renewable mouse intestinal epithelial cell monolayers and organoids for functional analyses. BMC Cell Biol. 2018 Aug 15;19(1):15. doi: 10.1186/s12860-018-0165-0.

[84] Kai Y. Intestinal villus structure contributes to even shedding of epithelial cells. Biophys J. 2021 Feb 16;120(4):699–710. doi: 10.1016/j.bpj.2021.01.003. Epub 2021 Jan 14.

[85] Acar E, Plopper GE, Yener B. Coupled analysis of in vitro and histology tissue samples to quantify structure-function relationship. PLoS One. 2012;7(3):e32227. doi: 10.1371/journal.pone.0032227. Epub 2012 Mar 30.

[86] Behroodi E, Latifi H, Najafi F. A compact LED-based projection microstereolithography for producing 3D microstructures. Sci Rep. 2019 Dec 23;9(1):19692. doi: 10.1038/s41598-019-56044-3. Erratum in: Sci Rep. 2020 Feb 17;10(1):2785. doi: 10.1038/s41598-020-59557-4.

[87] Jaiswal A, Rastogi CK, Rani S, Singh GP, Saxena S, Shukla S. Two decades of two-photon lithography: Materials science perspective for additive manufacturing of 2D/3D nano-microstructures. iScience. 2023 Mar 11;26(4):106374. doi: 10.1016/j.isci.2023.106374.

[88] Simmons DW, Schuftan DR, Ramahdita G, Huebsch N. Hydrogel-Assisted Double Molding Enables Rapid Replication of Stereolithographic 3D Prints for Engineered Tissue Design. ACS Appl Mater Interfaces. 2023 May 31;15(21):25313–25323. doi: 10.1021/acsami.3c02279. Epub 2023 May 18.

[89] Jinno S, Moeller HC, Chen CL, Rajalingam B, Chung BG, Dokmeci MR, Khademhosseini A. Microfabricated multilayer parylene-C stencils for the generation of patterned dynamic co-cultures. J Biomed Mater Res A. 2008 Jul;86(1):278–88. doi: 10.1002/jbm.a.32030.

[90] Hou L, Wang H, Zou H, Zhou Y. Robotic Manipulation Planning for Automatic Peeling of Glass Substrate Based on Online Learning Model Predictive Path Integral. Sensors (Basel). 2022 Feb 8;22(3):1292. doi: 10.3390/s22031292.

[91] Mizuno K, Hirashima T, Toda S. Robust tissue pattern formation by coupling morphogen signal and cell adhesion. EMBO Rep. 2024 Nov;25(11):4803–4826. doi: 10.1038/s44319-024-00261-z. Epub 2024 Sep 27.

[92] Sato T, Vries RG, Snippert HJ, van de Wetering M, Barker N, Stange DE, van Es JH, Abo A, Kujala P, Peters PJ, Clevers H. Single Lgr5 stem cells build crypt-villus structures in vitro without a mesenchymal niche. Nature. 2009 May 14;459(7244):262–5. doi: 10.1038/nature07935. Epub 2009 Mar 29.

[93] Heijmans J, van Lidth de Jeude JF, Koo BK, Rosekrans SL, Wielenga MC, van de Wetering M, Ferrante M, Lee AS, Onderwater JJ, Paton JC, Paton AW, Mommaas AM, Kodach LL, Hardwick JC, Hommes DW, Clevers H, Muncan V, van den Brink GR. ER stress causes rapid loss of intestinal epithelial stemness through activation of the unfolded protein response. Cell Rep. 2013 Apr 25;3(4):1128–39. doi: 10.1016/j.celrep.2013.02.031. Epub 2013 Mar 28.

[94] Schindelin J, Arganda-Carreras I, Frise E, Kaynig V, Longair M, Pietzsch T, Preibisch S, Rueden C, Saalfeld S, Schmid B, Tinevez JY, White DJ, Hartenstein V, Eliceiri K, Tomancak P, Cardona A. Fiji: an open-source platform for biological-image analysis. Nat Methods. 2012 Jun 28;9(7):676–82. doi: 10.1038/nmeth.2019.

[95] Staneva R, El Marjou F, Barbazan J, Krndija D, Richon S, Clark AG, Vignjevic DM. Cancer cells in the tumor core exhibit spatially coordinated migration patterns. J Cell Sci. 2019 Mar 15;132(6):jcs220277. doi: 10.1242/jcs.220277.

[96] Harris CR, Millman KJ, van der Walt SJ, Gommers R, Virtanen P, Cournapeau D, Wieser E, Taylor J, Berg S, Smith NJ, Kern R, Picus M, Hoyer S, van Kerkwijk MH, Brett M, Haldane A, Del Río JF, Wiebe M, Peterson P, Gérard-Marchant P, Sheppard K, Reddy T, Weckesser W, Abbasi H, Gohlke C, Oliphant TE. Array programming with NumPy. Nature. 2020 Sep;585(7825):357–362. doi: 10.1038/s41586-020-2649-2. Epub 2020 Sep 16.

[97] Hunter JD. Matplotlib: A 2D graphics environment. Comput Sci Eng. 2007;9:90–95. doi: 10.1109/MCSE.2007.55

[98] Legland D, Arganda-Carreras I, Andrey P. MorphoLibJ: integrated library and plugins for mathematical morphology with ImageJ. Bioinformatics. 2016 Nov 15;32(22):3532–3534. doi: 10.1093/bioinformatics/btw413. Epub 2016 Jul 13.

[99] Liberzon A, Käufer T, Bauer A, Vennemann P, Zimmer E. OpenPIV/openpiv-python: OpenPIV-Python v0.23.6 [software]. 2021. Available from: https://github.com/OpenPIV/openpiv-python

[100] Wang H, Barry K, Zaini A, Coakley G, Moyat M, Daunt CP, Wickramasinghe LC, Azzoni R, Chatzis R, Yumnam B, Camberis M, Le Gros G, Perdijk O, Foong JPP, Bornstein JC, Marsland BJ, Harris NL. Helminth infection driven gastrointestinal hypermotility is independent of eosinophils and mediated by alterations in smooth muscle instead of enteric neurons. PLoS Pathog. 2024 Aug 14;20(8):e1011766. doi: 10.1371/journal.ppat.1011766.

[101] Muraro D, Parker A, Vaux L, Filippi S, Almet AA, Fletcher AG, Watson AJM, Pin C, Maini PK, Byrne HM. Chronic TNF-driven injury delays cell migration to villi in the intestinal epithelium. J R Soc Interface. 2018 Aug;15(145):20180037. doi: 10.1098/rsif.2018.0037.

[102] Parker A, Vaux L, Patterson AM, Modasia A, Muraro D, Fletcher AG, Byrne HM, Maini PK, Watson AJM, Pin C. Elevated apoptosis impairs epithelial cell turnover and shortens villi in TNF-driven intestinal inflammation. Cell Death Dis. 2019 Feb 6;10(2):108. doi: 10.1038/s41419-018-1275-5.

[103] Vyhlidal CA, Chapron BD, Ahmed A, Singh V, Casini R, Shakhnovich V. Effect of Crohn’s Disease on Villous Length and CYP3A4 Expression in the Pediatric Small Intestine. Clin Transl Sci. 2021 Mar;14(2):729–736. doi: 10.1111/cts.12938. Epub 2020 Dec 16.

