## Supplemental Data for "Reconstituting organotypic 2D microtissue co-cultures via sequential stenciling"

Journal Name

Crossmark

RECEIVED
dd Month yyyy

REVISED
dd Month yyyy

PAPER

### Reconstituting organotypic 2D microtissue co-cultures via sequential stenciling

Kai Hirzel[1,2,*], Jasmin Cic[1,3,4,*], Stella Asmanidou[2], Nadja Schmohl[2], Roland E Kontermann[2,3], Satoshi Toda[5], Monilola A Olayioye[2,3], Andrew G Clark[2,3,4,†], Michael Heymann[1,3,†]

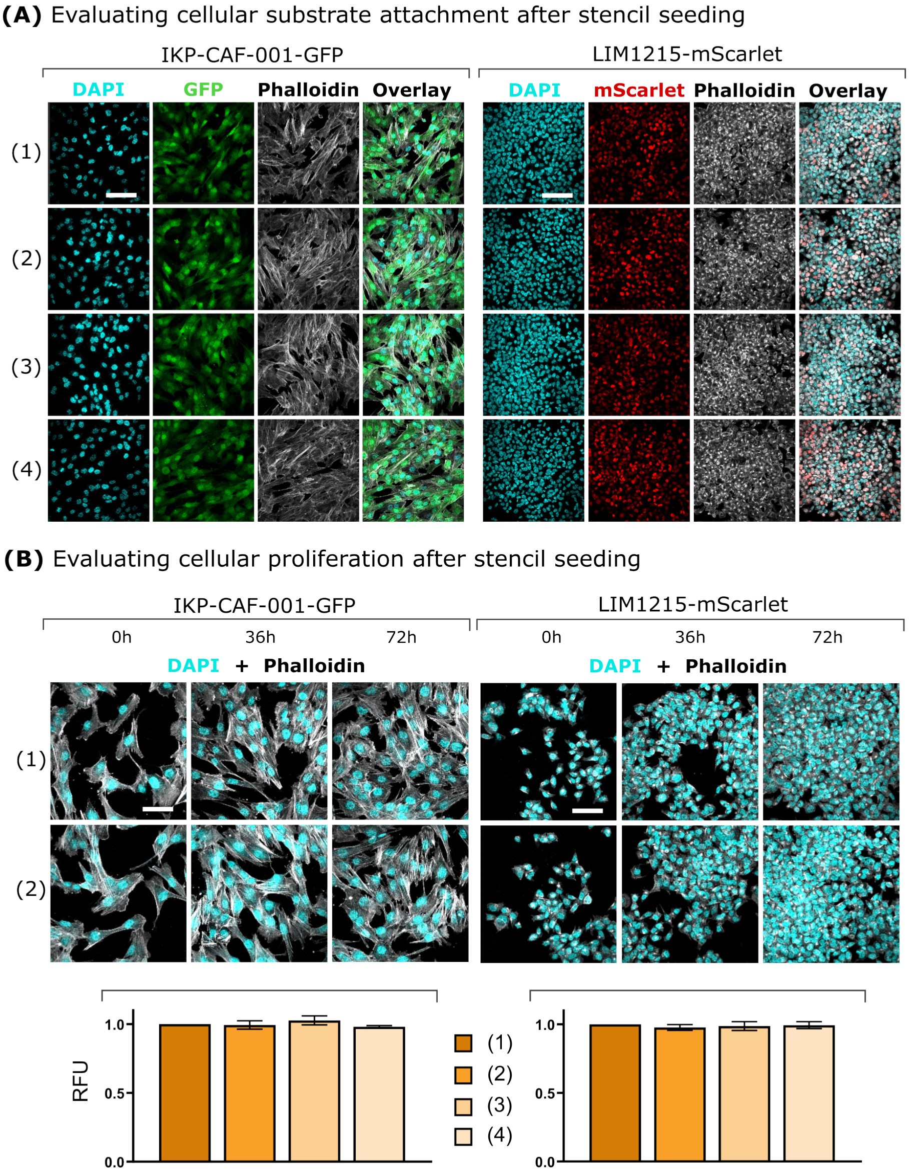


**SI Figure 1.** Evaluating cell-cultivation compatibility for cellular attachment, proliferation, and ATP metabolism. **(A)** Confocal immunofluorescence imaging for F-actin on fixed and stained IKP-CAF-001-GFP and LIM1215-Scarlet cells seeded either on a collagen-coated cover glass as a control (1), inside a 1 mm channel of a PDMS stencil (2), on the surface area of a removed stencil (3), or on the surface area of a removed stencil inside a 1 cm channel of a stencil (4) visualizes cell spreading on the substrate (scale bar 100 $\mu$m). **(B)** PDMS stencil influence on cell proliferation by quantifying cell confluency of fixed and stained cells using confocal immunofluorescence imaging (scale bar 50 $\mu$m). (C) ATP level measurement of IKP-CAF-001-GFP and LIM1215-Scarlet cells after trypsin-detachment from the substrate 24 hours after cell seeding (labels according to (A), n=3, N=30, error bars show the mean ± SD)

**SI Movies**

**SI Movie 1:** Time-lapse images of synNotch (GFP, mCherry and merged) gradient formation with rectangular co-culture patterns (left panels) and corresponding fluorescence intensity linescan quantification (right panel). Imaging was started 48 hours after stencil removal, and images were captured at 2h intervals for 72h. Selected timepoints are quantified in figure 6 in the main text. Scalebar 500 $\mu$m.

**SI Movie 2:** Time-lapse images of synNotch (GFP, mCherry and merged) gradient formation with circular co-culture patterns (top panels). For extraction of linescans, the circle pattern was straightened along the circumference of the pattern for both channels (bottom left panels). In the the corresponding fluorescence intensity linescan quantification (bottom right panel), position "0" refers to the center of the circular pattern. Imaging was started 48 h after stencil removal, and images were captured at 2 h intervals for 72 h. Selected timepoints are quantified in figure 6 in the main text. Scalebar 500 $\mu$m.

**SI Movie 3:** Time-lapse images of cell migration using brightfield (BF) imaging (top) and Particle Image Velocimetry (PIV) analysis (bottom) with single stencil patterning method. Images were captured at 30 min intervals for 12 h. Corresponding quantification averaged over time is shown in figure 8 in the main text. Scalebar 500 $\mu$m, Scale vector 1 $\mu$m/min.

**SI Movie 4:** Time-lapse images of cell migration using brightfield (BF) imaging (top) and Particle Image Velocimetry (PIV) analysis (bottom) with dual stencil patterning method. Images were captured at 30 min intervals for 12 h. Corresponding quantification averaged over time is shown in figure 8 in the main text. Scalebar 500 $\mu$m, Scale vector 1 $\mu$m/min.

**SI Design Files**

**SI design 1:** Extruded US AirForce target
**SI design 2:** Single cell circle $S_{\text{single}}$
**SI design 3:** Colorectal cancer $S_{\text{in}}$
**SI design 4:** Colorectal cancer $S_{\text{out}}$
**SI design 5:** synNotch $S_{\text{L}}$
**SI design 6:** synNotch $S_{\text{R}}$
**SI design 7:** Intestine $S_{\text{cv}}$
**SI design 8:** Intestine $S_{\text{block}}$
